# SOX4 suppresses vascular calcification via DNA-binding-independent regulation of the USP10–HNRNPC–WNT5A axis

**DOI:** 10.64898/2026.11.16.767198

**Authors:** Junbo Chen, He Cai, Xiaoyu Jin, Yi Li, Yingyi Zhu, Kefan Ma, He Liu, Xiaolong Zhu, Luyang Yu, Cong Qiu

## Abstract

**Background:** Vascular smooth muscle cell (VSMC) osteogenic differentiation drives vascular calcification, yet the endogenous mechanisms that restrain this transition remain poorly defined. SOX4 has recently emerged as a regulator of phenotype switching in endothelial and epithelial cells, but whether it governs VSMC fate and vascular calcification is unknown.

**Methods:** SOX4 expression was examined in human calcified arteries, in vitamin D3 (VitD3)-treated mouse aortas and in phosphate-treated human aortic smooth muscle cells. Loss- and gain-of-function experiments, an HMG-box-deleted SOX4 mutant, co-immunoprecipitation combined with mass spectrometry, ubiquitination and protein stability assays, RNA immunoprecipitation sequencing and mRNA decay assays were combined with VSMC-specific *Sox4* knockout mice and pharmacological WNT5A inhibition.

**Results:** SOX4 protein was upregulated in calcified VSMCs and arteries, yet SOX4 loss aggravated and SOX4 overexpression attenuated VSMC osteogenic differentiation. A SOX4 mutant lacking the HMG-box retained anti-calcific activity, placing the function outside DNA binding. SOX4 interacted with HNRNPC and attenuated the HNRNPC–USP10 association, promoting K48-linked ubiquitination and proteasomal degradation of HNRNPC. HNRNPC bound WNT5A mRNA and was required for its stability, so that SOX4 reduced WNT5A expression. VSMC-specific *Sox4* deletion aggravated VitD3-induced aortic calcification together with HNRNPC and WNT5A upregulation. Strengthening this braking axis with the WNT5A antagonist BOX5 attenuated VSMC osteogenic differentiation *in vitro* and vascular calcification *in vivo* in both genotypes, identifying WNT5A as a pharmacologically accessible node downstream of SOX4 loss.

**Conclusions:** SOX4 restrains VSMC osteogenic differentiation through post-transcriptional control of cell fate, defining a SOX4–HNRNPC–WNT5A axis and identifying it as a pharmacologically accessible target for vascular calcification.

## Introduction

Vascular calcification (VC), the pathological deposition of calcium-phosphate minerals in arterial walls, is a major independent risk factor for cardiovascular morbidity and mortality. VC is closely coupled to the osteogenic differentiation of vascular smooth muscle cells (VSMCs). Unlike most differentiated cells types, VSMCs exhibit remarkable phenotypic plasticity, allowing them to dedifferentiate and acquire characteristics of other cell lineages in response to changes in the local microenvironment.^1^ In pathological settings including chronic kidney disease, diabetes, and atherosclerosis, stimuli such as hyperphosphatemia and hypercalcemia trigger VSMCs to exhibit a profound phenotypic switch from a contractile, quiescent state to an osteogenic or chondrogenic-like phenotype.^2,3^ This transition involves increased expression of osteogenic markers, including runt-related transcription factor 2 (RUNX2), bone morphogenetic protein 2 (BMP2), and alkaline phosphatase (ALP), which collectively orchestrate the deposition of calcium-phosphate crystals within the extracellular matrix.^3–5^ Yet the endogenous mechanisms that restrain this transition remain poorly defined, which has limited the identification of nodes that can be reinforced therapeutically.

Transcription factors are major contributors to cell fate transition. Transcription factors are classically defined by their ability to bind DNA and regulate gene transcription. However, a growing body of evidence reveals that several transcription factors exert biological functions through DNA-binding-independent mechanisms, which have been shown to regulate protein stability or RNA metabolism by direct physical interaction with non-DNA partners.^6–8^ This non-canonical functional repertoire extends the regulatory scope of transcription factors beyond transcriptional control. However, the extent to which such non-canonical interactions operate in VSMCs remains largely unexplored, and elucidating these functions may offer new insights on the mechanisms of VSMC osteogenic differentiation.

SRY-related high-mobility group box 4 (SOX4) and its family member SOX9 have been increasingly recognized for their critical roles in cell fate determination such as epithelial-mesenchymal transition (EMT) and endothelial-mesenchymal transition (EndMT).^9–13^ Interestingly, the osteogenic differentiation of VSMCs involves reprogramming that shares features with both EMT and EndMT, including morphological change, extracellular matrix remodeling, and acquisition of a migratory phenotype.^1,4^ While the role of SOX9 in promoting VSMC osteogenic differentiation and vascular calcification has been well characterized,^14–16^ the specific function of SOX4 in this pathological process remains largely unknown. However, SOX4, which is unique among the SOXC factors, limits adult bone accrual by delaying osteoblastogenesis and promoting osteoclastogenesis in Lepr⁺ mesenchymal cells.^17^ In addition, SOX4 has been functionally linked to both the TGF-β and Wnt/β-catenin signaling pathways, which are closely associated with VSMC osteogenic differentiation and vascular calcification.^18–22^ These findings prompted us to determine whether SOX4 involves in VSMC osteogenic differentiation, and if so, through what mechanism.

Although SOX4 protein expression was significantly upregulated during the calcification process, its genetic loss in VSMCs markedly exacerbated osteogenic differentiation, suggesting that SOX4 acts as an endogenous suppressor of VSMC osteogenic differentiation. Mechanistically, SOX4 acts not through its canonical transcriptional activity, but by forming a complex with the RNA-binding protein HNRNPC, and attenuating the interaction between HNRNPC and ubiquitin-specific peptidase 10 (USP10), thereby promoting K48-linked ubiquitination and subsequent proteasomal degradation of HNRNPC. This process destabilizes *WNT5A* mRNA and suppresses its expression, thereby suppressing calcification. VSMC specific loss of SOX4 promotes vascular calcification, accompanied by increased HNRNPC and WNT5A expression. Enhancing the braking role of this regulatory axis using the WNT5A antagonist BOX5 markedly attenuates VSMC osteogenic differentiation and vascular calcification in both *in vitro* and *in vivo*. Our findings define a SOX4–HNRNPC–WNT5A pathway in VSMC osteogenic differentiation and identify it as a potential therapeutic target for intervention in vascular calcification.

## Methods

### Clinical samples

Protein lysates and formalin-fixed, paraffin-embedded tissue sections from human non-calcified and calcified arteries were kindly provided by the Fourth Affiliated Hospital of Zhejiang University School of Medicine. This retrospective study was approved by the hospital’s ethics committee with a waiver of informed consent and was conducted in accordance with the Declaration of Helsinki.

### Animal experiments

All procedures involving experimental animals received approval from the Animal Care and Research Committee of Zhejiang University (Approved code: ZJU20251096) and were conducted in accordance with the animal ethics guidelines established by the National Institutes of Health of the United States. Throughout the experiment, considerable attention was devoted to animal welfare, ensuring that both the number of experimental animals and their suffering were minimized. Mice were housed in the animal facilities at Zhejiang University in a humidity- and temperature-controlled environment (55 ± 5%, 22 ± 2 °C) with a 12 h light-dark cycle and no pathogenic microorganisms. *Sox4*^flox/flox^ mice were obtained from GemPharmatech Co., Ltd. Vascular smooth muscle cell-specific *Sox4* knockout mice were generated by crossing *Sox4*^flox/flox^ mice with *Myh11*^CreERT^^2^ mice to produce *Sox4*^flox/flox^/*Myh11*^CreERT2^ mice. Subsequent intraperitoneal injections of 75 mg/kg tamoxifen for 5 consecutive days were administered to the *Sox4*^flox/flox^/*Myh11*^CreERT2^ mice to generate *Sox4*^SMKO^ mice. *Sox4*^flox/flox^ mice without *Myh11*^CreERT2^ (*Sox4*^WT^) were referred to as control mice.

In order to induce vascular medial calcification, we administered 500,000 IU/kg of Vitamin D3 (VitD3, TargetMol, #T1135) subcutaneously to eight-week-old *Sox4*^WT^ and *Sox4*^SMKO^ mice over three consecutive days. The control group was treated with vehicle (corn oil). Tissue samples were harvested seven days post-treatment.

To investigate the therapeutic impact of BOX5 on vascular medial calcification, *Sox4*^WT^ and *Sox4*^SMKO^ mice were administered a daily intraperitoneal injection of 1 mg/kg of BOX5 (MCE, #HY-123071) for seven days following the subcutaneous injection of VitD3. The control group received saline as vehicle. Tissue samples were harvested seven days post-treatment.

All groups of mice were anaesthetized by isoflurane inhalation (3%) and euthanized by exsanguination (RWD, R510-22-10) before sample harvest. The entire aorta was dissected under an anatomical microscope for further analysis. Mice were randomly assigned to groups, and tissue collection and image quantification were performed by blinded, independent personnel.

### Tissue processing and frozen sectioning

The aortic arch was carefully dissected, fixed in 4% paraformaldehyde (PFA) for 15 minutes at room temperature, and dehydrated in 15% sucrose overnight at 4 °C. Tissues were then embedded in optimal cutting temperature (OCT) compound (SAKURA, #4583). Cryosections were cut at 5 μm thickness and stored at -80 °C for subsequent analyses. Before staining, sections were brought to room temperature, air-dried for 30 min, and rinsed with PBS for 3 times prior to use.

### Cell culture

Human aortic smooth muscle cells (hASMCs) were purchased from the American Type Culture Collection (ATCC) and cultured in high-glucose Dulbecco’s modified Eagle’s medium (DMEM, Gibco) containing 10% fetal bovine serum (FBS, Biochannel) and 1% penicillin/streptomycin (P/S, Gibco). Cells were cultured in a humidified incubator with 5% CO₂ at 37 °C, and passages 3-8 were used for all experiments to maintain VSMC phenotypic stability. To induce *in vitro* calcification, VSMCs were treated with calcification medium (culture medium containing 3 mM inorganic phosphate (#10039-32-4, Sangon Biotech)), with medium replacement every 2 days.

### Alizarin Red S and Von Kossa staining

To detect calcium deposition by calcium staining, the whole aorta or cultured hASMCs were fixed in 4% PFA at room temperature for 15 minutes. For Alizarin Red S staining, the entire aorta was incubated with 2% Alizarin Red S overnight at room temperature, while hASMCs in 12-well plates were incubated with 2% Alizarin Red S for 10 minutes at room temperature, followed by dual distilled water rinses. The aorta or hASMCs showing red/purple color were considered positive. For Von Kossa staining, vascular tissue sections were incubated with silver nitrate solution, exposed to ultraviolet light for 30 minutes, and rinsed with dual distilled water. The vessels showing black color were considered positive.

### Western blot

Aortic tissues were homogenized in radioimmunoprecipitation assay (RIPA) lysis buffer (Beyotime Biotechnology, #P0013C) supplemented with protease inhibitor cocktail (LABLEAD, C0101). For cell lysates, hASMCs were homogenized in a lysis buffer (50 mM Tris-HCl, 150 mM NaCl, 0.6% Triton X-100, 1 mM EDTA, 1 mM EGTA) containing protease inhibitors cocktail (LABLEAD, C0101) and phosphatase inhibitors cocktail (LABLEAD, C0104) to extract proteins. Subsequently, 50 μg of total protein underwent high-temperature denaturation, followed by electrophoresis and immediate transfer onto a PVDF membrane. The PVDF membrane was then blocked with TBST buffer containing 5% milk at room temperature for 1 hour. The membrane was incubated with primary antibodies overnight at 4 °C. After wash, the PVDF membrane was exposed to a secondary antibody conjugated with horseradish peroxidase at room temperature for 1 hour. Chemiluminescence detection was performed using an ECL kit (Yeasen Biotechnology, China, #36222). Protein band intensities were quantified with ImageJ and normalized to the corresponding GAPDH or β-actin levels. Details of the antibodies are provided in Table S1.

### Immunofluorescence (IF) staining

For frozen aortic sections, slides were brought to room temperature, rinsed with PBS, and blocked with 10% donkey serum, 1% BSA, and 0.3% Triton X-100 for 1 hour. Sections were incubated overnight at 4 °C with primary antibodies, washed three times with PBS, and then incubated with fluorescence-conjugated secondary antibodies and DAPI (1:1000) for 1 hour at room temperature. For paraffin sections, the slides were rehydrated and subjected to antigen retrieval according to the kit’s instructions (Beyotime, #P0088) before blocking and antibody incubation. For hASMCs, cells cultured on coverslips were fixed in 4% PFA for 15 minutes at room temperature, washed with PBS, and blocked with the same blocking solution for 1 hour. Cells were then incubated overnight at 4 °C with primary antibodies, followed by three washes with PBS and incubation with fluorescence-conjugated secondary antibodies and DAPI (1:1000) for 1 hour at room temperature. Stained sections and cells were examined using a fluorescence microscope (Olympus, FV3000), and fluorescence intensity was quantified using ImageJ. Details of the antibodies are listed in Table S1.

### Small interfering RNA (siRNA) transfection

siRNAs targeting the gene of *SOX4*, *HNRNPC*, *USP10* and negative control siRNAs (si-CTL) were synthesized by SUNYA Biotechnology (Hangzhou, China). hASMCs were seeded in 6-well plates at a density of 5 × 10⁵ cells/well and transfected with siRNA (final concentration: 50 nM) using Hieff Trans in vitro siRNA/miRNA Transfection Reagent (Yeasen Biotechnology, China, #40806) according to the manufacturer’s instructions. After 6 h of transfection, the medium was replaced with fresh complete medium, and cells were cultured for subsequent experiments (quantitative Real-Time PCR (qRT-PCR), Western blot, and calcification assays). The sequences of siRNA primers are listed in Table S2.

### Plasmid, Adenovirus and Lentivirus production

The plasmids of Flag-SOX4, HA-HNRNPC, and USP10 were purchased from UniBio (Chongqing, China). Adenoviruses carrying full-length SOX4 (Flag-SOX4-FL) and the HMG-box-deleted SOX4 mutant (Flag-SOX4-ΔHMG) were obtained from Hanheng Biotechnology (Shanghai, China). The lentivirus carrying HA-HNRNPC was purchased from Qingke Biology (Hangzhou, China).

### qRT-PCR

Total RNA was extracted from the hASMCs and mouse aortic tissues using the Super FastPure Cell RNA Isolation Kit (Vazyme, China, #RC102-01). Reverse transcription was performed using the HisyGo All-in-One RT Red SuperMix for qPCR (+gDNA Wiper) (Vazyme, China, #RT333-01), and mRNA levels were quantified via qRT-PCR using ChamQ Blue Universal SYBR qPCR Master Mix (Vazyme, China, #Q312-02) according to the manufacturer**’**s protocols. Relative expression levels were determined using the 2^−ΔΔCT^ method. All gene-specific primers used in this study are listed in the Table S3.

### Alkaline phosphatase (ALP) activity and quantification of hASMCs calcification

hASMCs underwent calcification induction by culturing them in a medium with 3 mM sodium phosphate buffer for 7 days, with the medium replenished every other day. Following this, the hASMCs were placed in cell lysate, centrifuged at 12000 rpm, and the resulting supernatant was collected. According to the manufacturer’s instructions of Alkaline Phosphatase Assay Kit (Beyotime, China, #P0322S), a standard curve was plotted with Para-nitrophenyl phosphate to determine the activity of ALP in the supernatant. For calcium assay, the Calcium Colorimetric Assay Kit (Beyotime, China, #S1063S) was utilized. Following the manufacturer’s instructions, a standard curve was established using calcium standard substances, and the calcium content in the supernatant was measured employing the O-cresol Phthalein complex one colorimetric method. The ALP activity and calcium content of each sample were normalized by the total protein concentration.

### Electrophoretic mobility shift (EMSA) assay

hASMCs were infected with Ad-Flag-SOX4-FL or Ad-Flag-SOX4-ΔHMG, and nuclear proteins were isolated 48 h later using a commercial nuclear and cytoplasmic protein extraction kit (FDbio, FD0199). EMSA was conducted using the LightShift Chemiluminescent EMSA Kit (Thermo, 20148) according to the manufacturer’s instructions. The sequences of the biotin-labeled probes is 5’-TCATAAGACAAAGAGAGACAAAGAGAGTGC-3’.

### Co-immunoprecipitation (Co-IP)

In Co-IP experiments, samples underwent homogenization in lysis buffer (50 mM Tris-HCl (pH 7.5), 150 mM NaCl, 1 mM EDTA, 1mM EGTA, 0.6% Triton X-100) supplemented with protease inhibitors. The protein extract was then incubated overnight at 4 °C with IgG and the primary antibody specific to the target protein. Subsequently, 40 μL protein A/G magnetic beads (MCE, HY-K0202) were introduced to each sample group and incubated at 4 °C with rotation for 2 hours. The captured protein samples were homogenized in 2x loading buffer containing protease inhibitors and phosphatase inhibitors before proceeding to Western Blot analysis. Details of the antibodies used for Co-IP are listed in Table S1.

### LC-MS/MS analysis

Anti-Flag antibodies were included in the lysate of hASMCs transfected with Flag-SOX4 for Co-IP, utilizing IgG as the negative control. LC-MS/MS analysis was conducted by Baipu Biotechnology Co., Ltd. (Shanghai, China). Substrate proteins potentially binding to SOX4 were identified through screening based on the scores of proteins detected in the mass spectrometry data and the Flag/IgG ratio.

### Protein stability assay

To assess protein stability, cells were first subjected to knockdown or overexpression as indicated, followed by treatment with cycloheximide (CHX, 50 μg/mL) or DMSO (vehicle control). At the indicated time points, cells were harvested and lysates were analyzed by Western blot. Target protein band intensities were quantified and normalized to GAPDH. The normalized values were then expressed relative to the 0 h time point.

### RIP assay, library preparation and sequencing

RIP experiments, high-throughput sequencing, and data analysis were performed by Seqhealth Technology Co., LTD (Wuhan, China). hASMCs (10 million) were washed twice with cold PBS and lysed in lysis buffer. 10% of the cell lysate was retained as “input” and 80% was subjected to immunoprecipitation using an anti-HNRNPC antibody (Proteintech, #68447-1-Ig). The remaining 10% was incubated with rabbit IgG (Abcam, #ab171870) as a negative control (“IgG”). The immunoprecipitation complexes were eluted after a series of washing steps. RNA was extracted using TRIzol reagent (Thermo Fisher, #15596018CN) according to the manufacturer’s instructions and quantified with a Qubit 3 fluorometer (Thermo Fisher). RNA integrity was assessed by agarose gel electrophoresis of the input RNA. Successful immunoprecipitation was validated by Western blotting. Libraries were prepared using the KCTM Digital Stranded mRNA Library Prep Kit (Kangce, Wuhan), which incorporates a 12-base unique molecular identifier (UMI) into pre-amplified cDNA molecules to correct for PCR and sequencing biases and errors. Library products containing 200 to 500 bp fragments were enriched and quantified, followed by sequencing on a DNBSEQ-T7 sequencer (MGI) in PE150 mode.

### RNA decay assay

hASMCs were seeded in six-well plates at a density of 5 × 10⁵ cells per well and allowed to adhere overnight. Cells were then subjected to the indicated knockdown or overexpression treatments. For the basal condition, following knockdown or overexpression, cells were treated with either DMSO (vehicle control) or actinomycin D (ActD, 5 μg/mL) and harvested at the indicated time points for RNA extraction. For the pathological condition, cells were first transfected with the indicated siRNAs or infected with adenoviruses/lentiviruses as described above; 24 h later, cells were switched to medium containing 3 mM Pi to induce osteogenic differentiation. After 3 days of Pi stimulation, ActD (5 μg/mL) or DMSO was added, and cells were harvested at the indicated time points. Total RNA was isolated using the Super FastPure Cell RNA Isolation Kit (Vazyme, China, #RC102-01) following the manufacturer’s protocol. The stability of *WNT5A* mRNA was assessed by qRT-PCR using ChamQ Blue Universal SYBR qPCR Master Mix (Vazyme, China, #Q312-02). The remaining mRNA levels at each time point were normalized to the 0-h time point (DMSO control) for each group.

### Cell viability assay

Cell viability was measured by cell-counting kit-8 (CCK8) assay (Beyotime Biotechnology, China, #C0038) according to the instruction protocols. VSMCs were treated with BOX5 at the indicated concentrations for the specified durations before viability assessment.

### Statistical analysis

Statistical analysis utilized GraphPad Prism 10.0 software (GraphPad Software). Data, presented as mean ± SD, were obtained from a minimum of three independent experiments. Prior to statistical analysis, the Shapiro-Wilke method was employed to assess normal distribution. Comparisons between the two groups were done using unpaired *t*-test. One-way analysis of variance (ANOVA) followed by Tukey’s multiple comparison test was used to compare the means among multiple groups. One-way ANOVA followed by Dunnett’s test was used for multiple comparisons against a control group. Two-way ANOVA analysis followed by a Bonferroni multiple comparison test was used for grouped analyses. Correlations in GTEx-derived human aortic expression data were assessed using Spearman correlation analysis. *P* < 0.05 was considered statistically significant.

### Data Availability

The RIP-seq data generated in this study have been deposited in the NCBI Sequence Read Archive (SRA) under the accession code “PRJNA1533086” (https://dataview.ncbi.nlm.nih.gov/object/PRJNA1533086?reviewer=oi8jlo45l5vvg6cbcbkc1jciv). The proteomic interaction data have been deposited in the ProteomeXchange Consortium via the iProX partner repository with the dataset identifier “PXD084684” (https://www.iprox.cn/page/PSV023.html;?url=1790224921281hxwd) (Password: D5Q9). All other relevant data are available from the corresponding author upon reasonable request.

## Results

### SOX4 is upregulated during VSMC osteogenic differentiation and in calcified arteries

To investigate the potential role of SOX4 in vascular calcification, we first examined SOX4 expression in human aortic smooth muscle cells (hASMCs) exposed to inorganic phosphate (Pi, 3 mM) to induce osteogenic differentiation *in vitro*.^23^ SOX4 was progressively upregulated from day 3 to day 7 during this process, accompanied by increased RUNX2 and BMP2, which are well-recognized markers of osteogenic differentiation (Figure 1A). Given the important role of VSMC osteogenic differentiation in vascular calcification, we further established a murine model of vascular calcification via subcutaneous injection of vitamin D3 (VitD3, 500,000 IU/kg/day) for three consecutive days.^24^ Immunoblotting showed elevated SOX4 protein levels in the aortas of VitD3-treated mice, along with increased RUNX2 and BMP2 (Figure 1B). Furthermore, immunofluorescence (IF) staining confirmed that SOX4 upregulation was mainly in vascular smooth muscle cells (VSMCs) within the calcified arterial wall (Figure 1C). These findings suggested a link between VSMC SOX4 expression and osteogenic differentiation in vascular calcification. To further validate these findings in human samples, we examined SOX4 expression in human calcified arteries. Immunoblotting of protein lysates from human calcified arteries revealed a marked upregulation of SOX4, along with increased RUNX2 and BMP2, compared with non-calcified controls (Figure 1D). Consistently, IF staining of human arterial sections showed that SOX4 expression was higher in calcified arteries than in non-calcified controls (Figure 1E). These data show that SOX4 expression is elevated in calcified VSMCs and arteries, and prompted us to investigate its functional role in VSMC osteogenic differentiation.

**Figure 1.**
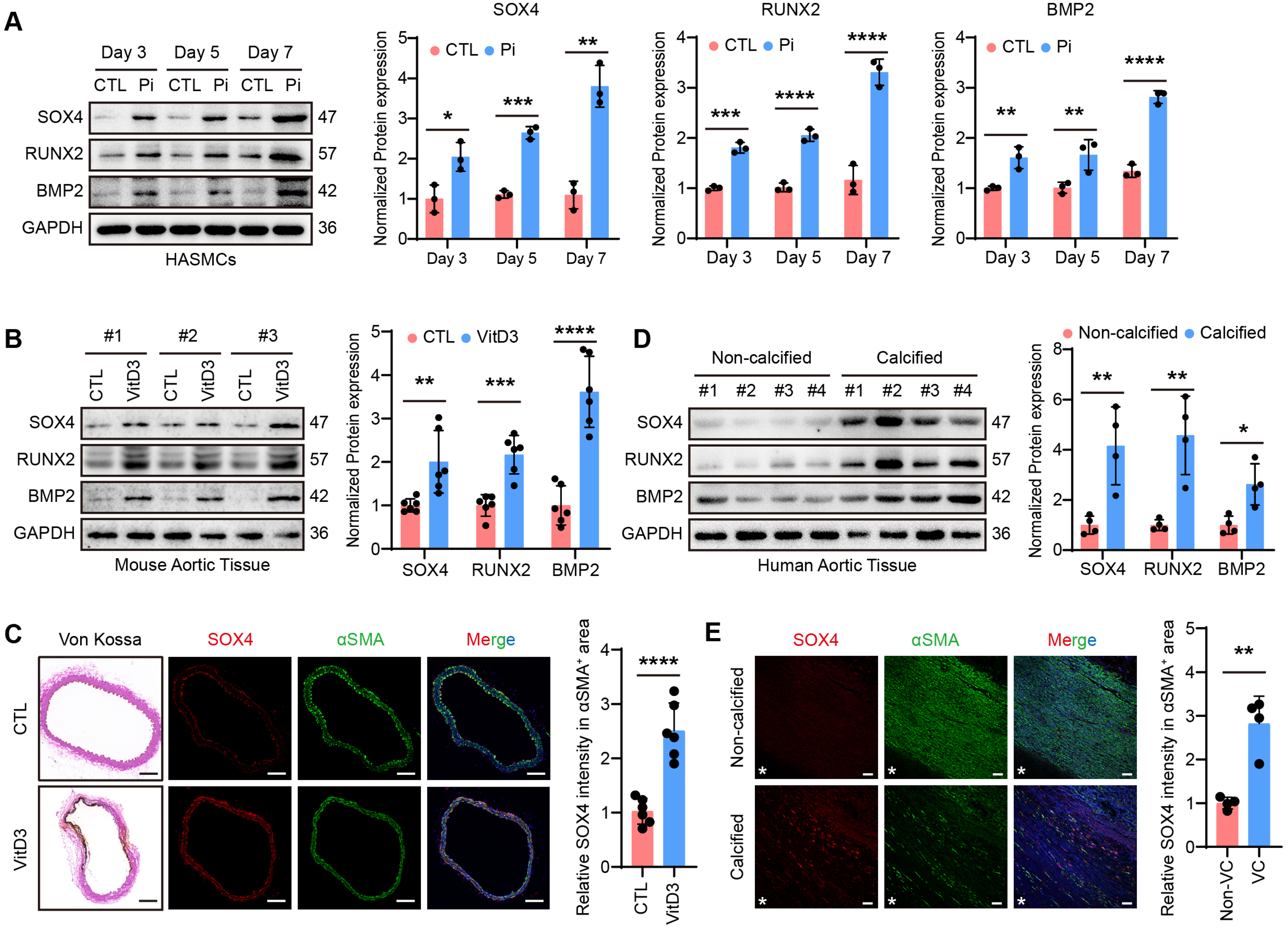
SOX4 expression is upregulated in calcified VSMCs and arteries. **A,** Immunoblot analysis and quantification of SOX4, RUNX2, and BMP2 in hASMCs treated with Pi (3 mM) for the indicated days. PBS was used as the vehicle control (n = 3). **B,** Immunoblot analysis and quantification of SOX4, RUNX2, and BMP2 in aortas from control (CTL, corn oil) and vitamin D3-treated (VitD3, 500,000 IU/kg/day) mice (n = 6). **C,** Representative Von Kossa staining and immunofluorescence images of SOX4 (red) and αSMA (green) in aortic sections from control and VitD3-treated mice, as described in (**B**), with quantification of relative SOX4 fluorescence intensity (n = 6). Scale bar: 100 μm. **D,** Immunoblot analysis and quantification of SOX4, RUNX2, and BMP2 in non-calcified and calcified human arteries (n = 4). **E,** Representative immunofluorescence images of SOX4 (red) and αSMA (green) in non-calcific and calcific human artery sections, with quantification of relative SOX4 fluorescence intensity (n = 4). Scale bar: 50 μm. Data are presented as mean ± SD and analyzed by unpaired *t*-test. \**P* < 0.05, \*\**P* < 0.01, \*\*\**P* < 0.001, \*\*\*\**P* < 0.0001.

### SOX4 suppresses VSMC osteogenic differentiation independent of its DNA-binding activity

To further explore the role of SOX4 in VSMC calcification, we employed loss- and gain-of-function approaches in cultured hASMCs. Transfection of SOX4-specific siRNA efficiently reduced SOX4 expression (Supplemental Figure S1A). SOX4 deficiency alone did not induce spontaneous calcification but significantly augmented Pi-induced VSMC calcification. Alizarin Red S staining revealed that SOX4-silenced cells exhibited markedly increased calcium nodule formation upon Pi stimulation (Figure 2A). Parallel to these changes, ALP activity and calcium deposition were also elevated in SOX4-deficient VSMCs upon Pi stimulation (Figure 2B, C), along with enhanced RUNX2 and BMP2 expression (Figure 2D). Conversely, SOX4 overexpression markedly suppressed Pi-induced VSMC osteogenic differentiation and calcification, as evidenced by reduced calcium nodule formation, diminished ALP activity, lower calcium deposition, and attenuated RUNX2 and BMP2 expression (Supplemental Figure S1B, Figure 2E–2H). These findings identify SOX4 as a negative regulator of VSMC calcification.

**Figure 2.**
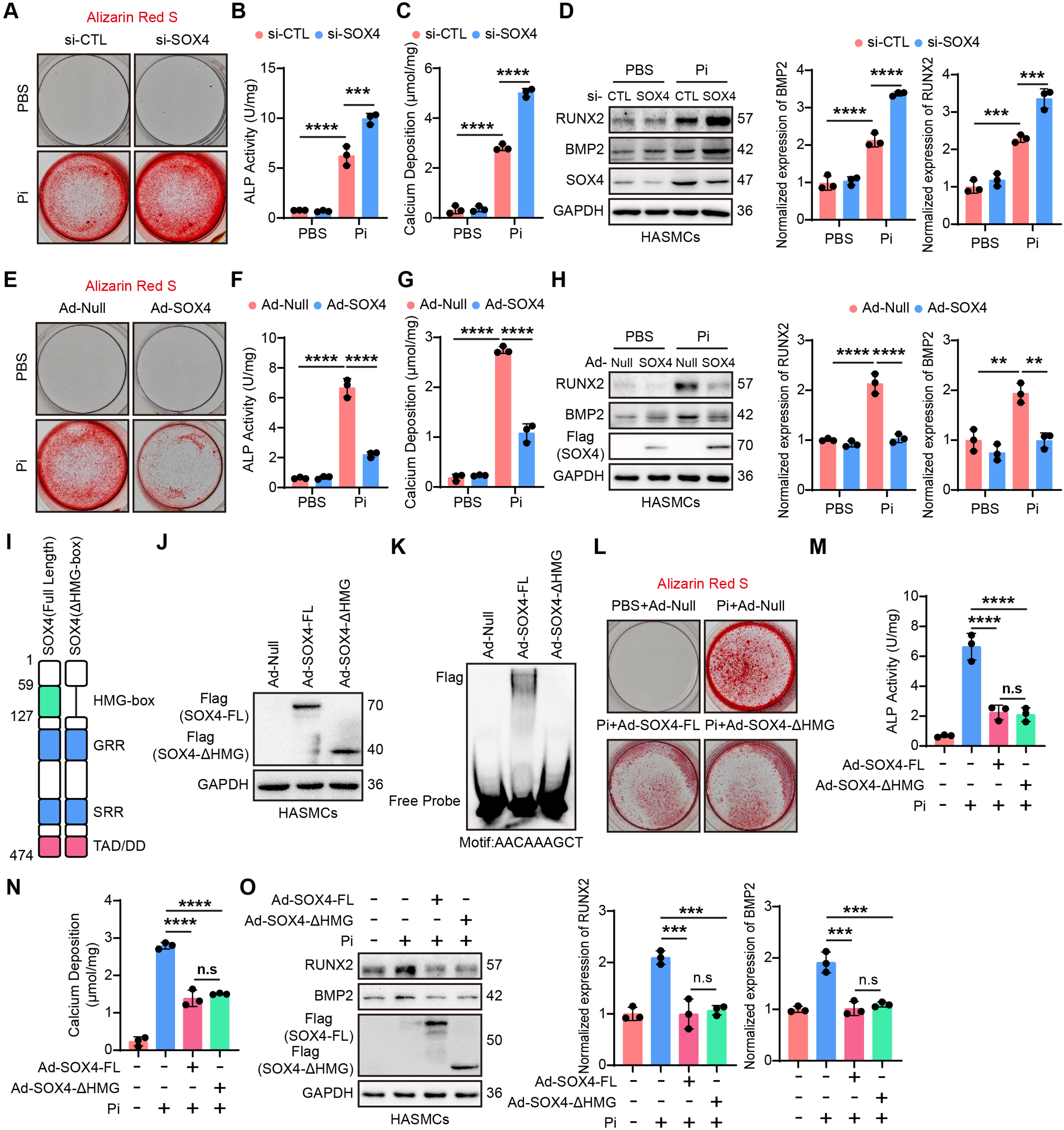
Deficiency of SOX4 aggravates VSMCs osteogenic differentiation independently of its DNA-binding. **A,** Representative images of Alizarin Red S staining of Pi (3 mM)-stimulated hASMCs pretreated with CTL siRNA or SOX4 siRNA (n = 3). **B, C,** ALP activity (**B**) and calcium deposition (**C**) in the indicated groups of cultured hASMCs (n = 3). **D,** Representative immunoblots and quantification of RUNX2 and BMP2 in hASMCs treated with vehicle (PBS) or Pi (3 mM) after transfection with control siRNA or SOX4 siRNA (n = 3). **E,** Representative Alizarin Red S staining of Pi (3 mM)-stimulated hASMCs pre-infected with Ad-Null or Ad-SOX4 (n = 3). **F, G,** ALP activity (**F**) and calcium deposition (**G**) in the indicated groups (n = 3). **H,** Representative immunoblots and quantification of RUNX2 and BMP2 in hASMCs treated with vehicle (PBS) or Pi (3 mM) after infection with Ad-Null or Ad-SOX4 (n = 3). **I,** Schematic illustration of the SOX4-ΔHMG-box construct. **J,** Representative immunoblots of Flag-SOX4-Full length (FL) and Flag-SOX4-ΔHMG in hASMCs infected with Ad-Null, Ad-SOX4-FL, or Ad-SOX4-ΔHMG. **K,** EMSA showing loss of DNA-binding activity by SOX4-ΔHMG. **L,** Representative images of Alizarin Red S staining in Pi (3 mM)-stimulated hASMCs pre-infected with ad-Null, Ad-SOX4-Full length (FL), or Ad-SOX4-ΔHMG (n = 3). **M, N,** ALP activity (**M**) and calcium deposition (**N**) in the indicated groups (n = 3). **O,** Representative immunoblots and quantification of RUNX2 and BMP2 in hASMCs treated with vehicle (PBS) or Pi (3 mM) after pre-infection with Ad-Null, Ad-SOX4-WT, or Ad-SOX4-ΔHMG (n = 3). Data are expressed as mean ± SD and analyzed by one-way ANOVA with Tukey’s test. \*\*\**P* < 0.001, \*\*\*\**P* < 0.0001. n.s, not significant.

To test whether the anti-calcific effect of SOX4 requires its DNA-binding activity, we generated a SOX4 mutant lacking the high-mobility group (HMG)-box DNA-binding domain (SOX4-ΔHMG) (Figure 2I, J). The HMG-box is a conserved DNA-binding motif that binds the minor groove, induces DNA bending, and recognizes the consensus motif A/TAACAA/T.^25^ EMSA confirmed that deletion of the HMG-box abolished SOX4 DNA-binding activity (Figure 2K). We therefore investigated the effect of SOX4-ΔHMG on VSMC osteogenic differentiation. The SOX4-ΔHMG mutant retained inhibitory activity against Pi-induced VSMC calcification comparable to that of full-length SOX4. Cells expressing SOX4-ΔHMG still exhibited reduced calcium nodule formation, decreased ALP activity and calcium deposition, as well as suppressed RUNX2 and BMP2 expression upon Pi stimulation (Figure 2L–2O). These results indicate that the HMG-box DNA-binding domain is dispensable for the anti-calcific function of SOX4, and that SOX4 restrains VSMC osteogenic differentiation by a mechanism that does not require its canonical DNA-binding activity.

### SOX4 interacts with HNRNPC and promotes its proteasomal degradation

To elucidate the molecular mechanism underlying SOX4-mediated inhibition of VSMC calcification, we sought to identify SOX4-interacting proteins. Co-immunoprecipitation (Co-IP) coupled with liquid chromatography-tandem mass spectrometry (LC-MS/MS) was performed in hASMCs. Intersection of the SOX4 interactome with Gene Ontology (GO) terms related to osteogenic differentiation identified two candidate proteins: HNRNPC and HSPE1 (Figure 3A). We prioritized HNRNPC because members of the HNRNP family are established regulators of smooth muscle cell differentiation.^26^ Moreover, accumulating evidence indicates that HNRNP proteins broadly govern stem cell lineage specification through both transcriptional and post-transcriptional mechanisms.^27^ However, whether HNRNPC contributes to VSMC osteogenic differentiation had not been tested. We therefore first determined its expression during this process. HNRNPC expression in hASMCs was upregulated during Pi-induced VSMC osteogenic differentiation (Figure 3B). This increase was recapitulated in the VitD3-induced mouse model, where HNRNPC was also markedly elevated in calcified aortas (Figure 3C, D). Consistently, immunoblotting of human arterial samples showed that HNRNPC protein levels were higher in calcified arteries than in non-calcified controls (Figure 3E). These results indicate that HNRNPC is associated with osteogenic differentiation in vascular calcification.

**Figure 3.**
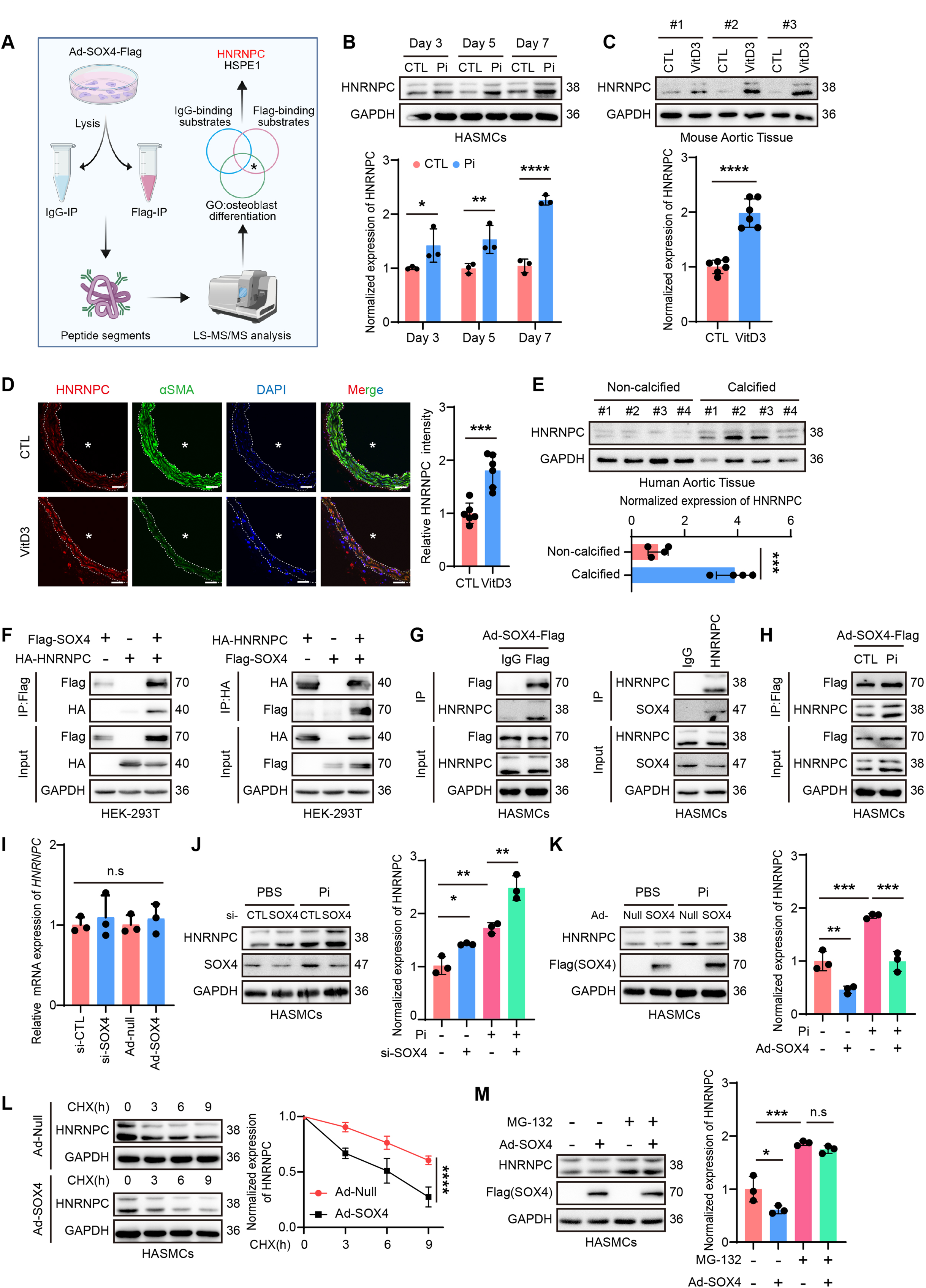
SOX4 interacts with HNRNPC and promotes its proteasomal degradation. **A,** Schematic illustration of mass spectrometry analysis of the SOX4 protein complex. HNRNPC was identified as a SOX4-interacting protein associated with osteogenic differentiation. **B,** Representative immunoblots and quantification of HNRNPC in hASMCs treated with Pi (3 mM) for the indicated days. PBS was used as the vehicle control (n = 3). **C,** Representative immunoblots and quantification of HNRNPC levels in aortic lysates from control (CTL, corn oil) or VitD3 (500,000 IU/kg/day)-treated mice (n = 6). **D,** Representative immunofluorescence images of HNRNPC (red) and αSMA (green) in aortas from control (CTL, corn oil) and vitamin D3-treated (VitD3, 500,000 IU/kg/day) mice, with quantification of relative HNRNPC intensity (n = 6). White asterisks indicate the lumen. Scale bar: 100 μm. **E,** Representative immunoblots and quantification of HNRNPC in non-calcified and calcified human arteries (n = 4). **F,** Representative Co-IP immunoblots of SOX4 and HNRNPC interaction in 293T cells. **G,** Representative Co-IP blots of endogenous SOX4 and HNRNPC interaction in hASMCs. **H,** Representative Co-IP blots of SOX4 and HNRNPC interaction in hASMCs following Pi (3 mM) stimulation for 3 days. **I,** Relative mRNA level of *HNRNPC* in hASMCs that treated as indicated (n = 3). **J,** Representative immunoblots and quantification of HNRNPC in hASMCs treated with vehicle (PBS) or Pi (3 mM) after transfection with control siRNA or SOX4 siRNA (n = 3). **K,** Representative immunoblots and quantification of HNRNPC in hASMCs treated with vehicle (PBS) or Pi (3 mM) after pre-infection with Ad-Null or Ad-SOX4 (n = 3). **L,** Representative immunoblots and quantification of HNRNPC in VSMCs pre-infected with Ad-Null or Ad-SOX4 and treated with CHX (50 μg/mL) for the indicated time points. The HNRNPC/GAPDH ratio was normalized to the zero-time point (DMSO) of each group (n = 3). **M,** Representative immunoblots and quantification of HNRNPC in VSMCs pre-infected with Ad-Null or Ad-SOX4 and treated with MG-132 (50 μg/mL) for 6 h before harvest. The HNRNPC/GAPDH ratio was normalized to the Ad-Null + DMSO group (n = 3). Data are expressed as mean ± SD. Unpaired *t*-test was used in (**B, C, D,** and **E)**; One-way ANOVA with Tukey’s test was used in (**I**-**K, M**); Two-way ANOVA with Bonferroni’s test was used in (**l**). \**P* < 0.05, \*\**P* < 0.01, \*\*\**P* < 0.001, \*\*\*\**P* < 0.0001. n.s, not significant.

To investigate how SOX4 regulates HNRNPC, we first investigated their association. Co-IP showed the HNRNPC–SOX4 interaction in HEK293T cells expressing Flag-SOX4 and HA-HNRNPC, and at the endogenous level in hASMCs (Figure 3F, G). Consistent with the HMG-box-independent anti-calcific effect of SOX4, HNRNPC also co-immunoprecipitated with the SOX4-ΔHMG mutant at both exogenous and endogenous levels (Supplemental Figure S2), indicating that the SOX4–HNRNPC interaction does not require the HMG-box. Under Pi stimulation, the HNRNPC–SOX4 interaction was enhanced compared with basal conditions, suggesting its involvement in osteogenic differentiation (Figure 3H).

Having established their association, we next asked whether SOX4 regulates HNRNPC expression. Neither SOX4 knockdown nor overexpression affected *HNRNPC* mRNA levels (Figure 3I). However, SOX4 knockdown in hASMCs increased HNRNPC protein levels, whereas SOX4 overexpression reduced them, especially under calcifying conditions (Figure 3J, K). In 293T cells, SOX4 overexpression also decreased HNRNPC expression (Supplemental Figure S3A). These results demonstrate that SOX4 negatively regulates HNRNPC protein expression under both normal and pathological conditions. We therefore investigated whether SOX4 alters HNRNPC protein stability. Cycloheximide chase assays revealed that SOX4 overexpression accelerated HNRNPC protein degradation in both hASMCs and 293T cells (Figure 3L, Supplemental Figure S3B). Given that HNRNPC has been reported to undergo ubiquitin-proteasomal degradation,^28^ we investigated whether proteasome inhibition with MG-132 could reverse the downregulation of HNRNPC caused by SOX4 overexpression. As expected, MG-132 treatment abolished SOX4-mediated HNRNPC downregulation (Figure 3M). These data indicate that SOX4 interacts with HNRNPC and promotes its proteasomal degradation.

### SOX4 attenuates the USP10–HNRNPC interaction to promote HNRNPC degradation

How does SOX4 promote HNRNPC degradation? We first tested whether the effect is ubiquitin-dependent. In Co-IP assays, loss of SOX4 was accompanied by reduced HNRNPC polyubiquitination, whereas SOX4 overexpression increased it (Figure 4A–4C). Using ubiquitin mutants restricted to single linkage types, we found that the SOX4-dependent increase was specific to K48-linked ubiquitination of HNRNPC (Figure 4D). To identify the deubiquitinase opposing this modification, we examined co-expression in human aortic tissue using the GTEx database, which showed a positive correlation between USP10 and HNRNPC transcript levels (Supplemental Figure S4A). Because co-expression does not establish a functional pairing, we tested the relationship experimentally. Knockdown of USP10 in VSMCs reduced HNRNPC protein levels, whereas USP10 overexpression in 293T cells increased them (Supplemental Figure S4B, C). Furthermore, Co-IP assays confirmed the HNRNPC–USP10 interaction in hASMCs (Figure 4E) and in 293T cells overexpressing both proteins (Figure 4F). Given that USP10 is a deubiquitinase known to regulate protein ubiquitination in vascular smooth muscle cells,^29,30^ we then examined whether USP10 regulates HNRNPC ubiquitination. Loss of USP10 increased HNRNPC polyubiquitination and reduced its expression in VSMCs, whereas USP10 overexpression had the opposite effect (Supplemental Figure S4D, E). These data indicate that USP10 stabilizes HNRNPC by reducing its ubiquitination.

**Figure 4.**
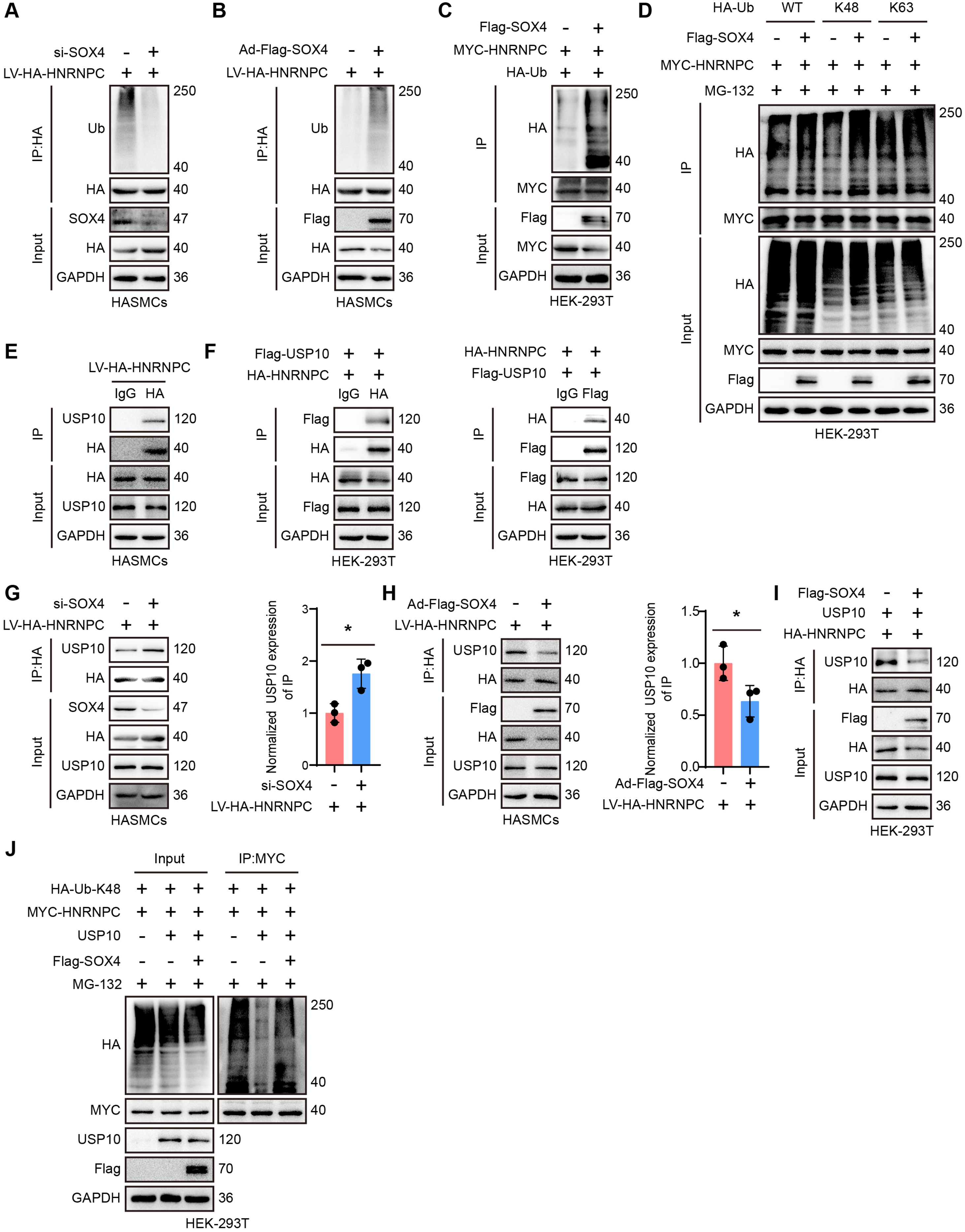
SOX4 promotes HNRNPC ubiquitination by suppressing the HNRNPC-USP10 interaction. **A, B,** Representative immunoblots of HNRNPC ubiquitination in hASMCs after SOX4 knockdown (**A**) or SOX4 overexpression (**B**). **C,** Representative blots of HNRNPC ubiquitination in 293T cells co-transfected with HNRNPC and ubiquitin, with or without SOX4 overexpression. **D,** Co-IP analysis of HNRNPC ubiquitination in 293T cells co-expressing HNRNPC with wildtype (WT), K48-only or K63-only ubiquitin, with or without SOX4 co-expression. Cells were treated with MG-132 (50 μg/mL) for 6 h before harvest. **E,** Representative Co-IP blots of endogenous HNRNPC–USP10 interaction in hASMCs. **F,** Representative Co-IP blots of HNRNPC–USP10 interaction in 293T cells. **G, H,** Representative Co-IP blots and quantification of HNRNPC–USP10 interaction in hASMCs after SOX4 knockdown (**G**) or SOX4 overexpression (**H**). **I,** Representative Co-IP blots of HNRNPC–USP10 interaction in 293T cells co-transfected with HNRNPC and USP10, with or without SOX4 overexpression. **J,** Representative Co-IP blots of HNRNPC ubiquitination in 293T cells co-expressing HNRNPC and K48-only ubiquitin, with or without USP10 and SOX4 as indicated. Cells were treated with MG-132 (50 μg/mL) for 6 h before harvest. Representative blots are from three independent experiments. The quantification data are expressed as mean ± SD and analyzed by Unpaired *t*-test. \**P* < 0.05.

We then asked whether SOX4 affects the USP10–HNRNPC association. Of note, SOX4 knockdown enhanced this interaction while SOX4 overexpression attenuated it in hASMCs (Figure 4G, H). Moreover, in 293T cells, SOX4 overexpression also attenuated their binding (Figure 4I). These findings raise the possibility that SOX4 restrains VSMC osteogenic differentiation by limiting the USP10–HNRNPC association. Consistent with this, in 293T cells USP10 expression reduced HNRNPC K48-linked ubiquitination, and additional SOX4 co-expression attenuated this effect (Figure 4J).

Together, these findings suggest that SOX4 inhibits the USP10–HNRNPC interaction, leading to HNRNPC polyubiquitination and degradation, and thereby suppressing VSMC osteogenic differentiation.

### SOX4 Inhibits VSMC osteogenic differentiation in an HNRNPC-dependent manner

To determine whether SOX4 inhibits VSMC calcification in an HNRNPC-dependent manner, we performed rescue experiments using HNRNPC-specific siRNA and overexpression constructs. Knockdown of HNRNPC in SOX4-silenced hASMCs significantly attenuated the exacerbated calcification caused by SOX4 deficiency, as evidenced by reduced calcium nodule formation, decreased ALP activity, calcium deposition, and diminished RUNX2 and BMP2 expression (Figure 5A–5E). Conversely, HNRNPC overexpression reversed the protective effects of SOX4 overexpression against Pi-induced VSMC calcification. In cells co-overexpressing both SOX4 and HNRNPC, the reduced calcium nodule formation, decreased ALP activity, and diminished calcium deposition observed with SOX4 overexpression alone were all restored to levels comparable to control cells (Figure 5F–5J). Accordingly, the suppression of RUNX2 and BMP2 expression by SOX4 was also largely abolished upon HNRNPC co-overexpression.

**Figure 5.**
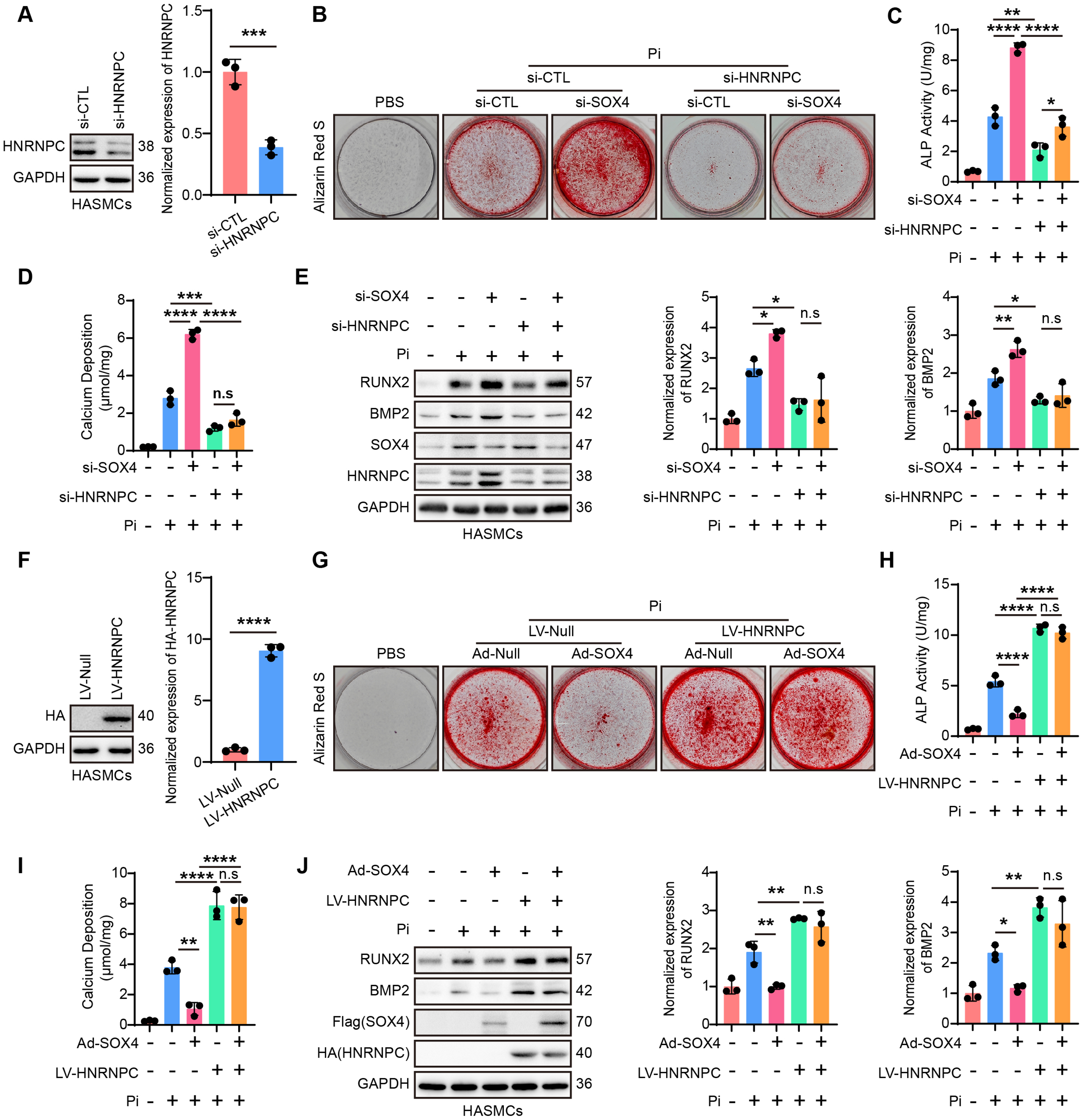
SOX4 inhibits VSMC calcification in an HNRNPC-dependent manner. **A,** Representative immunoblots and quantification of HNRNPC following HNRNPC siRNA treatment (n = 3). **B,** Representative images of Alizarin Red S staining in Pi (3 mM)-stimulated hASMCs transfected with CTL siRNA, SOX4 siRNA, or SOX4 siRNA plus HNRNPC siRNA as indicated (n = 3). **C, D,** ALP activity (**C**) and calcium deposition (**D**) in hASMCs treated as in (**B**) (n = 3). **E,** Representative immunoblots and quantification of RUNX2 and BMP2 in hASMCs treated as in (**B**) (n = 3). **F,** Representative immunoblot and quantification of HNRNPC following LV-HNRNPC infection (n = 3). **G,** Representative images of Alizarin Red S staining in Pi (3 mM)-stimulated hASMCs pre-infected with Ad-SOX4 or Ad-SOX4 plus LV-HNRNPC as indicated (n = 3). **H, I** ALP activity (**H**) and calcium deposition (**I**) in hASMCs treated as in (**G**) (n = 3). **J,** Representative immunoblots and quantification of RUNX2 and BMP2 in hASMCs treated as in (**G**) (n = 3). Data are expressed as mean ± SD. Unpaired *t*-test was used in (**A**) and (**F**); One-way ANOVA with Tukey’s test was used in (**C-E**) and (**H-J**). \**P* < 0.05, \*\**P* < 0.01, \*\*\**P* < 0.001, \*\*\*\**P* < 0.0001. n.s, not significant.

These loss- and gain-of-function experiments indicate that HNRNPC is required for the anti-calcific effect of SOX4 and acts downstream of it. The ability of HNRNPC overexpression to counteract SOX4 overexpression further indicates that HNRNPC abundance is critical for VSMC osteogenic differentiation under these conditions. This finding aligns with and reinforces our observation that SOX4 exerts its inhibitory function by reducing HNRNPC protein levels through accelerated proteasomal degradation. Thus, HNRNPC serves as an essential downstream effector through which SOX4 regulates VSMC osteogenic differentiation and calcification.

### SOX4 regulates *WNT5A* mRNA stability via HNRNPC

As a member of the heterogeneous nuclear ribonucleoprotein family, HNRNPC is involved in multiple aspects of RNA metabolism, including RNA stability,^28,31,32^ alternative splicing,^33–35^ and recognition of m6A modifications.^36^ Yet the precise mechanisms by which it exerts this function are unclear. To identify calcification-associated transcripts associated with HNRNPC in VSMCs, we conducted RNA immunoprecipitation sequencing (RIP-seq) (Supplemental Figure S5). This analysis identified 4,121 HNRNPC-bound RNAs, predominantly mRNAs enriched in coding sequence (CDS) regions (Figure 6A). GO enrichment analysis revealed that HNRNPC-associated mRNAs were significantly enriched in osteogenic differentiation-related pathways (Figure 6B). Among these, WNT5A, a known promoter of vascular calcification via non-canonical Wnt signaling,^37^ ranked as the most abundantly enriched transcript within this category (Figure 6C, Table S4).

**Figure 6.**
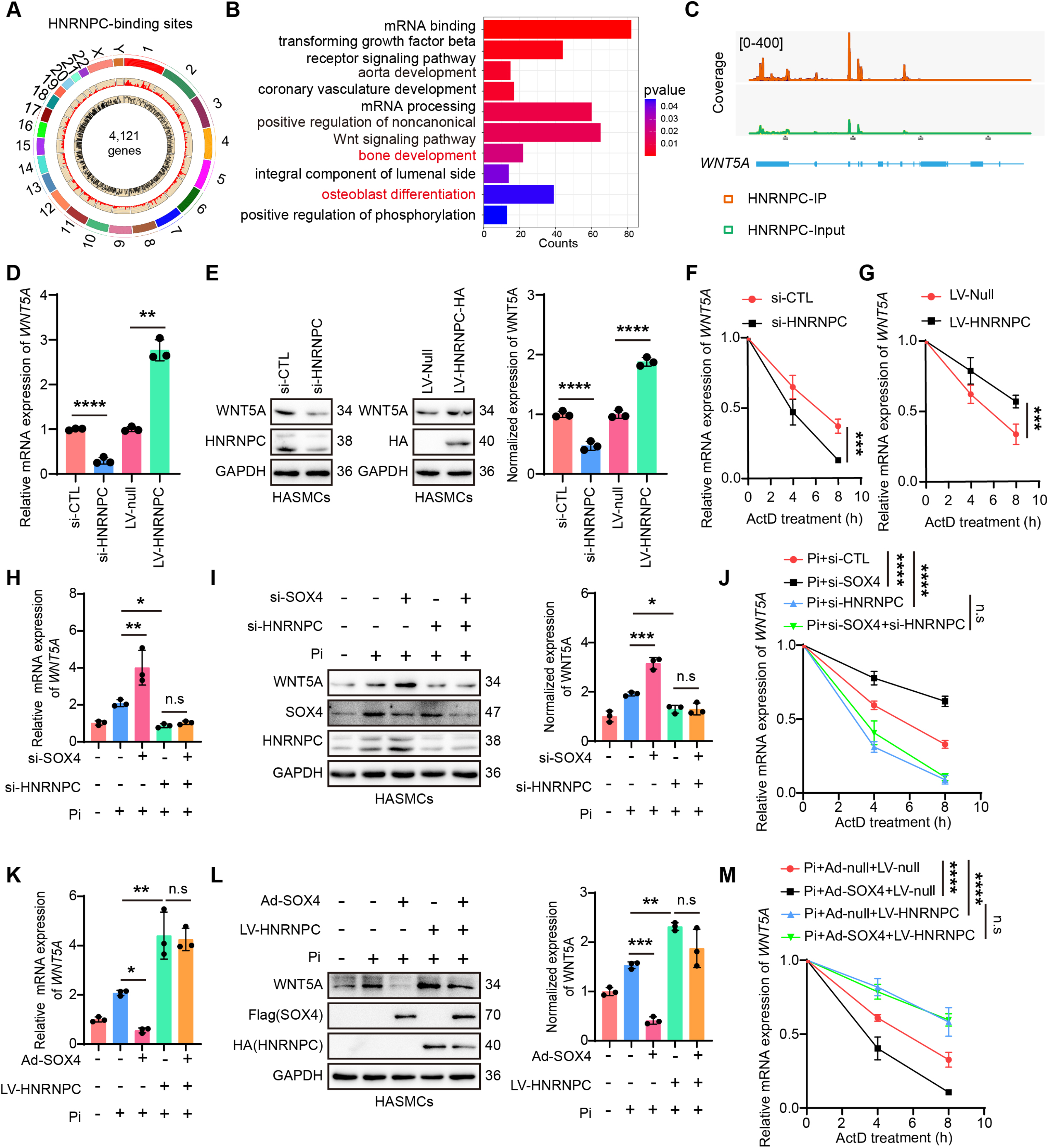
SOX4 regulates *WNT5A* mRNA stability via HNRNPC. **A,** Circular plot showing the HNRNPC-binding sites identified by RIP-seq analysis in hASMCs. **B,** Gene Ontology (GO) term analysis of 4121 genes bound by HNRNPC in hASMCs. **C,** Integrative Genomics Viewer tracks showing the HNRNPC-binding peaks at the *WNT5A* locus in the RIP-seq data. **D,** Relative mRNA levels of *WNT5A* in hASMCs following HNRNPC knockdown or overexpression, as indicated (n = 3). **E,** Representative immunoblots and quantification of WNT5A in hASMCs following HNRNPC knockdown or overexpression, as indicated (n = 3). **F,** Relative mRNA levels of *WNT5A* in hASMCs pretreated with CTL siRNA or HNRNPC siRNA, followed by ActD (5 µg/mL) for the indicated times. mRNA levels were normalized to the 0-h time point of each group (n = 3). **G,** Relative mRNA levels of *WNT5A* in hASMCs pretreated with LV-Null or LV-HNRNPC, followed by ActD (5 µg/mL) for the indicated times. mRNA levels were normalized to the 0-h time point of each group (n = 3). **H,** Relative mRNA levels of *WNT5A* in hASMCs treated with SOX4 siRNA, HNRNPC siRNA, or SOX4 plus HNRNPC siRNAs under Pi stimulation (3 mM, 3 days) (n = 3). **I,** Representative immunoblots of WNT5A in hASMCs treated with SOX4 siRNA, HNRNPC siRNA, or SOX4 plus HNRNPC siRNAs under Pi stimulation (3 mM, 3 days) (n = 3). **J,** Relative mRNA levels of *WNT5A* in hASMCs treated with SOX4 siRNA, HNRNPC siRNA, or SOX4 plus HNRNPC siRNAs under Pi stimulation (3 mM, 3days), followed by ActD (5 µg/mL) for the indicated times. mRNA levels were normalized to the 0-h time point of each group (n = 3). **K,** Relative mRNA levels of *WNT5A* in hASMCs pre-infected with Ad-SOX4, LV-HNRNPC, or Ad-SOX4 plus LV-HNRNPC under Pi stimulation (3 mM, 3 days) (n = 3). **L,** Representative immunoblots of WNT5A in hASMCs pre-infected with Ad-SOX4, LV-HNRNPC, or Ad-SOX4 plus LV-HNRNPC under Pi stimulation (3 mM, 3 days) (n = 3). **M,** Relative mRNA levels of *WNT5A* in hASMCs pre-infected with Ad-SOX4, LV-HNRNPC, or Ad-SOX4 plus LV-HNRNPC under Pi stimulation (3 mM, 3 days), followed by ActD (5 µg/mL) for the indicated times. mRNA levels were normalized to the 0-h time point of each group (n = 3). Data are expressed as mean ± SD. Unpaired *t*-test was used in (**D**) and (**E**); One-way ANOVA with Tukey’s test was used in (**H, I, K,** and **l**); Two-way ANOVA with Bonferroni’s test was used in (**F, G, J,** and **M**). \**P* < 0.05, \*\**P* < 0.01, \*\*\**P* < 0.001, \*\*\*\**P* < 0.0001. n.s, not significant.

We first asked whether HNRNPC controls WNT5A output. Under basal conditions, HNRNPC knockdown decreased WNT5A mRNA and protein levels, while overexpression had the opposite effect (Figure 6D, E). Importantly, HNRNPC knockdown abolished the upregulation of WNT5A caused by SOX4 deficiency (Supplemental Figure S6A, B), suggesting that SOX4 acts through HNRNPC to regulate WNT5A. Because HNRNPC functions in RNA metabolism, we tested whether this control operates at the level of transcript stability. Actinomycin D chase assays showed that HNRNPC knockdown accelerated WNT5A mRNA decay, whereas overexpression prolonged its half-life (Figure 6F, G). Moreover, depleting HNRNPC reversed the increase in WNT5A stability induced by SOX4 deficiency (Supplemental Figure S6C).

Whether SOX4 regulates WNT5A mRNA stability through HNRNPC under pathological conditions was then examined. Upon Pi stimulation, HNRNPC silencing reversed the increases in WNT5A mRNA and protein levels and the enhanced mRNA stability caused by SOX4 deficiency (Figure 6H–6J). Conversely, HNRNPC overexpression counteracted the decreases in WNT5A mRNA and protein levels and the reduced mRNA stability induced by SOX4 overexpression (Figure 6K–6M). These findings indicate that SOX4, acting through HNRNPC, reduces the stability and abundance of WNT5A mRNA to restrain VSMC osteogenic differentiation.

### VSMC-specific deletion of SOX4 aggravates vascular calcification *in vivo*

To validate the functional relevance of the SOX4–HNRNPC–WNT5A axis *in vivo*, we generated inducible VSMC-specific *Sox4* knockout mice (*Sox4*^SMKO^) by crossing *Sox4*-floxed mice with *Myh11*-CreERT2 transgenic mice. *Sox4*-floxed mice lacking *Myh11*-CreERT2 served as wild-type controls (*Sox4*^WT^) (Supplemental Figure S7A–S7B). Six-week-old *Sox4*^SMKO^ and *Sox4*^WT^ mice received intraperitoneal injections of tamoxifen (75 mg/kg) for five consecutive days to induce Cre-mediated recombination, resulting in efficient SOX4 deletion in aortic tissues (Supplemental Figure S7C–S7E). At eight weeks of age, mice were subcutaneously injected with either vehicle (corn oil) or VitD3 for three days, and aortas were harvested seven days later (Figure 7A).

**Figure 7.**
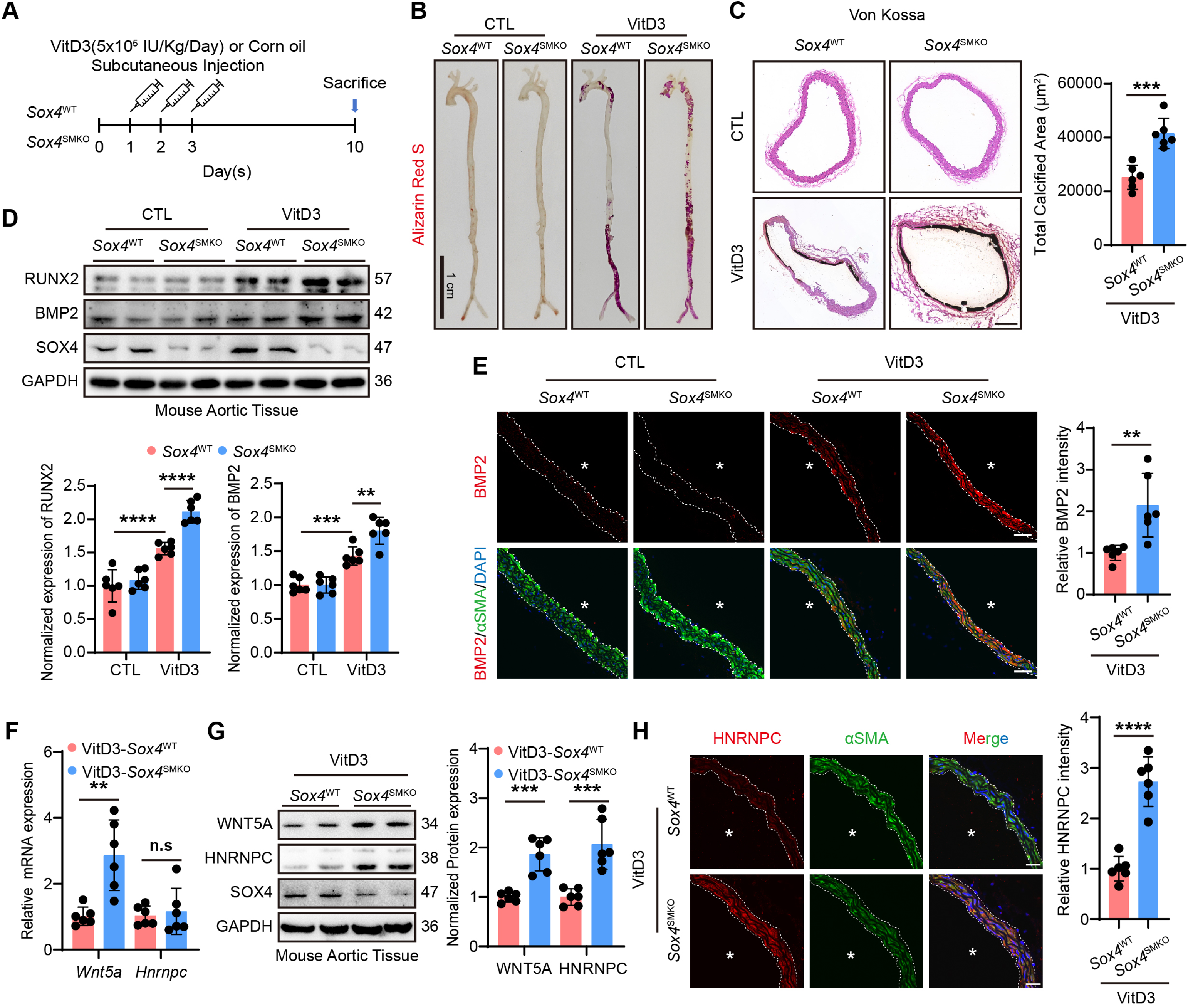
Deficiency of SOX4 aggravates vascular calcification. **A,** Schematic illustration of vascular calcification induced by VitD3 (500,000 IU/kg/day) in two groups of mice. **B,** Representative images of Alizarin Red S staining from whole aortas in two groups of mice that received vehicle (CTL, corn oil) or VitD3 injection (n = 6). Scale bar: 1 cm. **C,** Representative Von Kossa staining of aortic calcification in control (CTL, corn oil) and VitD3-treated mice, with quantification of total calcified area (n = 6). Scale bar: 100 μm. **D,** Representative immunoblots and quantification of RUNX2 and BMP2 in aortas from control (CTL, corn oil) and VitD3-treated mice (n = 6). **E,** Representative immunofluorescence images of BMP2 (red) and αSMA (green) from the aortic section in two groups of mice that received vehicle (CTL) or VitD3, with quantification of relative BMP2 intensity (n = 6). White asterisks indicate the lumen. Scale bar: 25 μm. **F,** Relative mRNA levels of *Wnt5a* and *Hnrnpc* in aortas from VitD3-treated *Sox4*^WT^ and *Sox4*^SMKO^ mice (n = 6). **G,** Representative immunoblots and quantification of HNRNPC and WNT5A in aortas from VitD3-treated *Sox4*^WT^ and *Sox4*^SMKO^ mice (n = 6). **H,** Representative immunofluorescence images of HNRNPC (red) and αSMA (green) in aortas from VitD3-treated *Sox4*^WT^ and *Sox4*^SMKO^ mice, with quantification of relative HNRNPC intensity (n = 6). White asterisks indicate the lumen. Scale bar: 25 μm. Data are expressed as mean ± SD. Unpaired *t*-test was used in (**C, E, F**-**H**); One-way ANOVA with Tukey’s test was used in (**D**). \*\**P* < 0.01, \*\*\**P* < 0.001, \*\*\*\**P* < 0.0001. n.s, not significant.

Alizarin Red S staining of whole aortas showed that SOX4 deficiency alone did not trigger spontaneous calcification, but significantly aggravated VitD3-induced aortic calcification compared with *Sox4*^WT^ mice (Figure 7B). Von Kossa staining of aortic cross-sections also revealed no baseline differences in medial wall thickness or calcium deposition between *Sox4*^WT^ and *Sox4*^SMKO^ mice. However, following VitD3 challenge, *Sox4*^SMKO^ mice exhibited reduced medial wall thickness and substantially increased calcium deposition (Figure 7C). Immunoblotting analysis showed elevated RUNX2 and BMP2 protein levels in the aortas of *Sox4*^SMKO^ mice following VitD3 treatment (Figure 7D). Consistent with this, IF staining further confirmed enhanced BMP2 expression in VSMC of *Sox4*-deficient arteries upon VitD3 challenge (Figure 7E). These results demonstrate that VSMC-specific *Sox4* deletion promotes vascular calcification and VSMC osteogenic differentiation.

We next examined whether the SOX4–HNRNPC–WNT5A axis was altered *in vivo*. In the VitD3-induced calcification model, qRT-PCR and immunoblotting of aortic lysates showed that both WNT5A mRNA and protein levels were significantly elevated in *Sox4*^SMKO^ mice compared with *Sox4*^WT^ controls (Figure 7F, G). Consistent with our *in vitro* findings, VSMC SOX4 deficiency had no effect on HNRNPC mRNA levels but markedly upregulated HNRNPC protein levels (Figure 7F–7H). These findings show that SOX4 deficiency aggravates vascular calcification in association with increased HNRNPC protein, but not mRNA, and increased WNT5A mRNA and protein, consistent with the axis defined *in vitro*.

### Pharmacological inhibition of WNT5A with BOX5 attenuates vascular calcification

To confirm that SOX4 suppresses VSMC calcification through the HNRNPC–WNT5A axis, we tested BOX5, a specific WNT5A antagonist, and asked whether it could reverse the exacerbated calcification caused by SOX4 deficiency (Figure 8A).^38–40^ We first examined its cytotoxicity in VSMCs using the CCK-8 assay. BOX5 showed no cytotoxicity at concentrations up to 100 µM (Figure 8B), which was used for all subsequent experiments. *In vitro*, BOX5 treatment significantly attenuated Pi-induced VSMC calcification in both control and SOX4-silenced cells. BOX5 reduced calcium nodule formation, ALP activity, calcium deposition, and RUNX2/BMP2 expression in SOX4-deficient cells to levels comparable to those in control cells, effectively abolishing the pro-calcific effect of SOX4 deficiency (Figure 8C–8F). These *in vitro* results show that BOX5 suppresses Pi-induced VSMC osteogenic differentiation and attenuates the exacerbated calcification resulting from SOX4 deficiency, indicating that WNT5A signaling is required for the pro-calcific effect of SOX4 loss.

**Figure 8.**
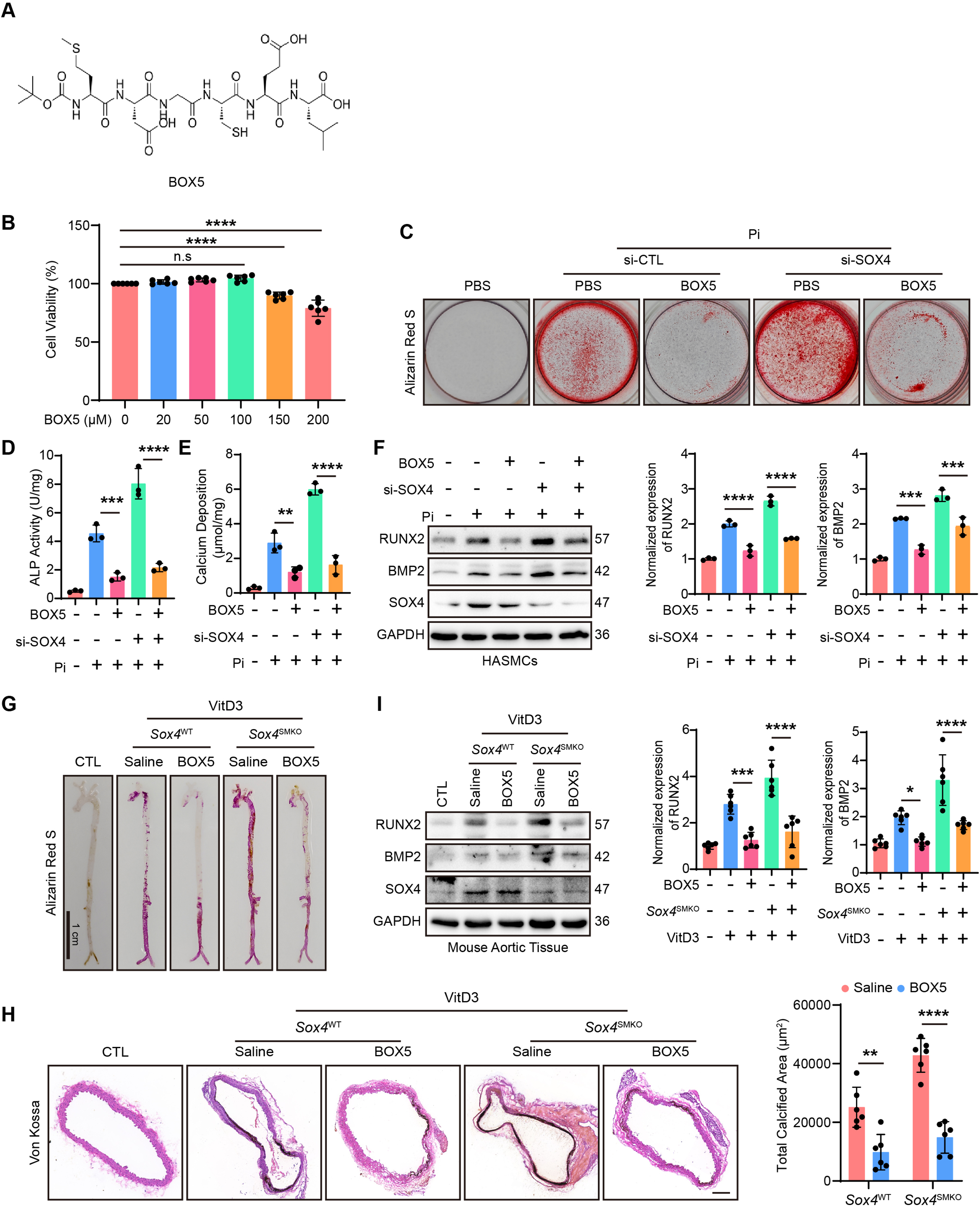
Pharmacological inhibition of WNT5A with BOX5 attenuates VSMC osteogenic differentiation and vascular calcification induced by SOX4 deficiency. **A**, Chemical structure of BOX5. **B,** Cell viability of hASMCs treated with increasing concentrations of BOX5 (n = 6). **C,** Representative images of Alizarin Red S staining from hASMCs. After SOX4 knockdown, cells were incubated with Pi (3 mM) or vehicle (PBS) for 3 days and then exposed to BOX5 (100 μM) or vehicle (PBS) for additional 3 days before staining (n = 3). **D, E,** ALP activity (**D**) and calcium deposition (**E**) in hASMCs treated as in (**C**) (n = 3). **F,** Representative immunoblots and quantification of RUNX2 and BMP2 in hASMCs treated as in (**C**) (n = 3). **G,** Representative images of Alizarin Red S staining of whole aortas from *Sox4*^WT^ and *Sox4*^SMKO^ mice. After *Sox4* knockout in smooth muscle cells, mice were treated with VitD3 (500,000 IU/kg/day) for 3 days and then exposed to BOX5 (1 mg/kg/day) or saline for consecutive 7 days before harvest (n = 6). Control (CTL) mice were *Sox4*^WT^ mice receiving corn oil as vehicle. Scale bar: 1 cm. **H,** Representative images of Von Kossa staining and quantification of aortic calcification in the five groups treated as in (**G**) (n = 6). Scale bar: 100 μm. **I,** Representative immunoblots and quantification of RUNX2 and BMP2 in aortas from the five groups treated as in (**G**) (n = 6). Data are expressed as mean ± SD. One-way ANOVA with Dunnett’s test was used in (**B**); One-way ANOVA with Tukey’s test was used in (**D**-**F, H,** and **I**). \**P* < 0.05, \*\**P* < 0.01, \*\*\**P* < 0.001, \*\*\*\**P* < 0.0001. n.s, not significant.

To assess the *in vivo* efficacy of BOX5, *Sox4*^WT^ and *Sox4*^SMKO^ mice were subjected to VitD3-induced calcification and received daily intraperitoneal injections of BOX5 (1 mg/kg/day) for one week.^39,41^ Alizarin Red S and Von Kossa staining revealed that BOX5 administration markedly attenuated aortic calcification in both genotypes. Notably, BOX5 treatment not only reduced the calcified area in *Sox4*^SMKO^ mice to levels comparable to those observed in vehicle-treated *Sox4*^WT^ controls, effectively abolishing the exacerbated calcification phenotype resulting from SOX4 deficiency, but also attenuated VitD3-induced calcification in *Sox4*^WT^ mice, bringing it to a similar baseline level (Figure 8G, H). Consistently, the RUNX2 and BMP2 expression levels in aortic tissues of both genotypes were reduced. The elevated expression levels in *Sox4*^SMKO^ mice were reduced to levels comparable to vehicle-treated controls, while those in VitD3-treated *Sox4*^WT^ mice were also restored to baseline (Figure 8I).

Collectively, these data indicate that the SOX4–HNRNPC–WNT5A axis serves as an endogenous inhibitory mechanism against vascular calcification, yet its basal activity is insufficient to fully prevent calcification. Pharmacological reinforcement of this axis with the WNT5A antagonist BOX5 reduced calcification in control mice and reversed the exacerbated calcification in *Sox4*^SMKO^ mice. These findings establish WNT5A as a key mediator of SOX4-dependent protection and support the SOX4–HNRNPC–WNT5A axis as a potential therapeutic target for vascular calcification.

## Discussion

Vascular calcification (VC), characterized by pathological calcium-phosphate deposition in arterial walls, is a significant and independent risk factor for cardiovascular morbidity and mortality.^42^ It is highly prevalent in patients with chronic kidney disease (CKD), diabetes, and atherosclerosis, contributing to increased arterial stiffness, reduced vascular compliance, and ultimately, adverse cardiovascular events such as myocardial infarction and stroke.^43,44^ Elucidating the molecular mechanisms governing this pathological process is therefore critical for the development of targeted interventions. The central finding of this study is that SOX4 acts as an endogenous suppressor of VSMC osteogenic differentiation through a mechanism that does not require HMG-box-mediated DNA binding. Instead, it interacts with the RNA-binding protein HNRNPC and disrupts the HNRNPC–USP10 association, thereby promoting K48-linked ubiquitination and proteasomal degradation of HNRNPC. The resulting reduction in HNRNPC abundance destabilizes *WNT5A* mRNA, lowering WNT5A expression and restraining the osteogenic program in VSMCs. *In vivo*, VSMC-specific deletion of SOX4 aggravated VitD3-induced aortic calcification, and pharmacological inhibition of WNT5A with BOX5 reversed this phenotype (Figure 9). Thus, the SOX4–HNRNPC–WNT5A axis defines a non-canonical regulatory module governing VSMC osteogenic differentiation, with vascular calcification as a downstream consequence.

**Figure 9.**
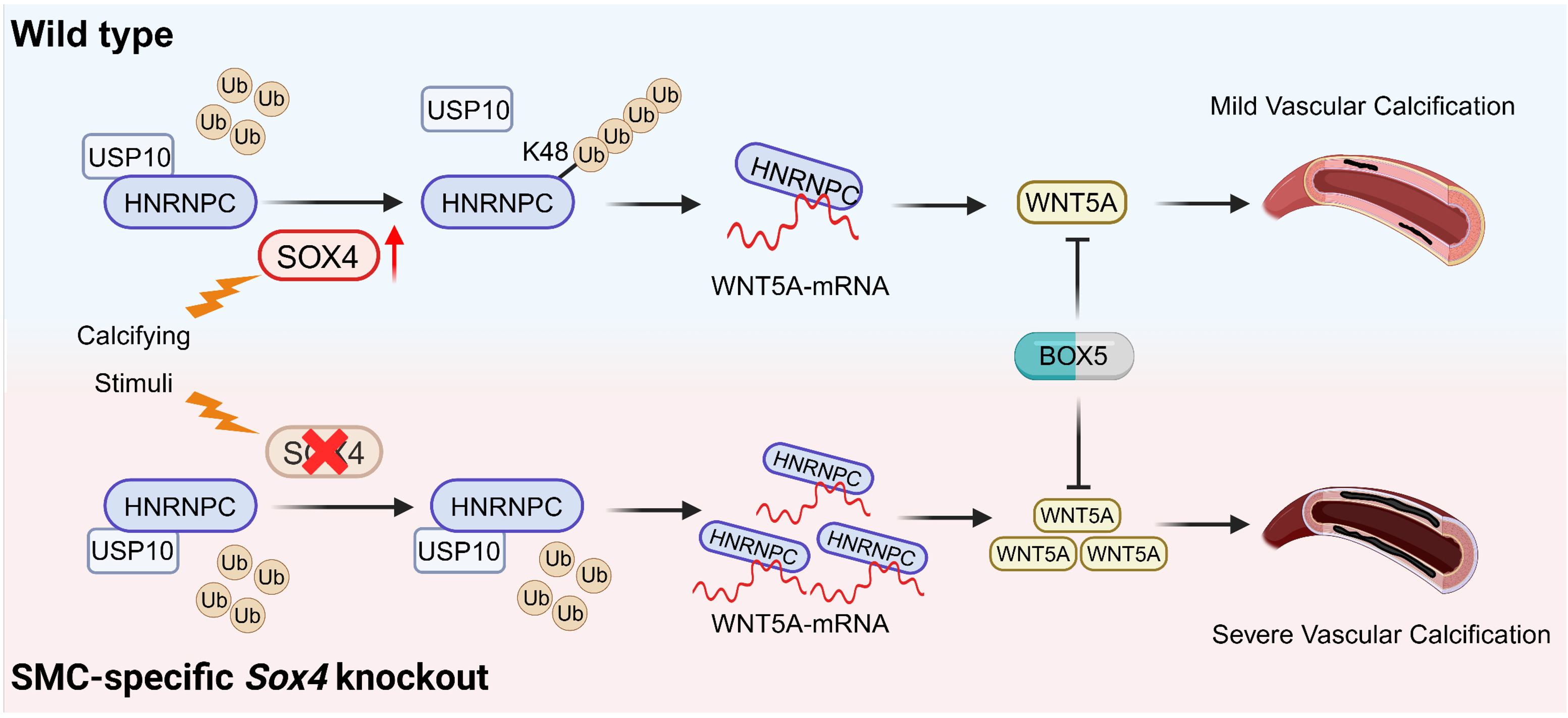
Schematic illustration of SOX4 suppresses vascular calcification by destabilizing *WNT5A* via regulation of USP10-HNRNPC interaction. SOX4 acts as an endogenous suppressor of arterial calcification by directly interacting with the RNA-binding protein HNRNPC, and disrupting the HNRNPC–USP10 interaction, thereby promoting K48-linked ubiquitination and proteasomal degradation of HNRNPC. This process destabilizes *WNT5A* mRNA and suppresses its expression, consequently attenuating calcification. Pharmacological inhibition of WNT5A with its specific antagonist BOX5 effectively mitigates VSMC osteogenic differentiation and vascular calcification.

SOX4 belongs to the SOXC family and contains a conserved HMG-box DNA-binding domain. In pathological contexts, SOX4 has been implicated primarily as a disease-promoting factor. For instance, SOX4 expression is aberrantly elevated in numerous cancers, where it accelerates tumor progression by promoting EMT and fostering an immunosuppressive microenvironment.^9^ In the cardiovascular system, SOX4 has also been identified as a pathogenic driver of atherosclerosis, cardiac hypertrophy, and myocardial ischemia.^10,45,46^ In contrast, our study shows that SOX4 deficiency accelerates VSMC osteogenic differentiation and vascular calcification despite its elevated expression, revealing a protective role in this context. This discrepancy may reflect cell type-specific functions of SOX4 among distinct cardiovascular cell populations, as its role in VSMCs may differ from that in cardiomyocytes, endothelial cells, or macrophages. Of note, this protective role also contrasts with that of SOX9, a related SOX family member known to promote VSMC osteogenic differentiation and vascular calcification.^14–16^ The opposing functions of SOX4 and SOX9 raise the possibility of a balance between pro- and anti-calcific SOX activities in the diseased vessel wall, although we did not test this directly. The paradoxical upregulation of protective factors during disease progression has been documented previously. For instance, NAMPT expression is elevated in calcified human and murine arteries, yet its ablation exacerbates VitD3-induced vascular calcification.^47^ Similarly, Ucma/GRP (Gla-rich protein) is markedly upregulated in calcifying VSMCs in response to elevated phosphate, whereas Ucma/GRP-deficient VSMCs exhibit enhanced osteogenic differentiation and calcification.^48^ These observations support the concept that SOX4 upregulation in calcified VSMCs may represent an adaptive, compensatory feedback mechanism that functions as a brake to limit disease progression. Notably, while SOX4 protein levels were elevated upon pro-calcific stimulation, its mRNA levels remained unchanged (Supplemental Figure S8A–S8C). Interestingly, SOX4 is regulated by USP10 in T cells,^49^ and we confirmed the SOX4-USP10 interaction in VSMCs (Supplemental Figure S8D). These observations suggest that altered USP10-mediated polyubiquitination may contribute to SOX4 upregulation in VSMCs under pathological conditions. However, whether additional post-translational modifications or chaperone-mediated stabilization also participate, and how this regulation integrates with the SOX4–USP10–HNRNPC axis described above, remain to be determined.

The HMG-box domain of SOX4 confers sequence-specific DNA-binding activity, enabling it to activate or repress target gene transcription. Unexpectedly, a SOX4 mutant lacking the HMG-box domain retained anti-calcific activity, comparable to full-length SOX4. This finding demonstrates that SOX4’s anti-calcific function is uncoupled from its HMG-box-mediated DNA binding. However, we note a distinction that our data support only in part. The ΔHMG mutant is DNA-binding-deficient by EMSA and retains anti-calcific activity, which establishes that HMG-box-mediated DNA binding is dispensable for this function. It does not formally exclude a residual contribution from transcriptional regulation, for two reasons: deletion of the HMG-box removes a protein-interaction surface as well as the DNA-contacting element, and we did not compare the transcriptomes of cells expressing full-length and ΔHMG SOX4 or determine their subcellular distributions. A DNA-binding-deficient mutant that preserves the domain fold, together with comparative transcriptomics, would be needed to close this gap. We therefore describe the mechanism as HMG-box-mediated DNA-binding independent throughout, and reserve “transcription-independent” for the possibility that remains open.

Through mass spectrometry-based interactome analysis, we identified HNRNPC as a key SOX4-associated protein. SOX4 binds to HNRNPC and promotes its ubiquitin-proteasomal degradation, thereby suppressing VSMC osteogenic differentiation and vascular calcification. Thus, SOX4 operates through a non-canonical, protein–protein interaction-dependent mechanism in this context. This non-transcriptional function of SOX4 has also been reported. For example, SOX4 binds p53 and interferes with MDM2-mediated ubiquitination, thereby stabilizing p53 protein.^50^ Here, we identified USP10 as a critical deubiquitinase that controls HNRNPC ubiquitination and subsequent proteasomal degradation. We further demonstrated that SOX4 inhibits the USP10–HNRNPC interaction, thereby promoting HNRNPC degradation and reducing its expression. Nevertheless, two aspects of the ubiquitin-dependent step remain incompletely defined. First, we show that USP10 stabilizes HNRNPC and that SOX4 weakens the USP10– HNRNPC association, but we did not test a catalytically inactive USP10 mutant. We therefore cannot exclude that USP10 acts in part as a steric shield that limits access of the ubiquitination machinery to HNRNPC, although the reciprocal effects of USP10 knockdown and overexpression on HNRNPC polyubiquitination are consistent with an enzymatic contribution. Second, the E3 ligase that catalyses K48-linked ubiquitination of HNRNPC is not identified here. Our data fit a model in which SOX4 raises HNRNPC ubiquitination by limiting USP10-dependent editing of ongoing ligase activity, but they do not exclude an alternative in which SOX4 recruits an E3 ligase to HNRNPC. These models make different predictions about the composition of the SOX4– HNRNPC complex and would require reconstitution with purified components to distinguish. We accordingly describe SOX4 as a modulator of ubiquitin-dependent HNRNPC turnover rather than as a defined subunit of a specific ligase complex. Given that SOX4 also binds USP10 (Supplemental Figure S8D), it may exert a dual regulatory function: disrupting the USP10– HNRNPC interaction to promote HNRNPC ubiquitination, while simultaneously engaging USP10 to limit its own ubiquitination and enhance its stability. This model, although speculative, would explain how SOX4 protein accumulates under calcifying conditions without a corresponding increase in mRNA. Further structural and biochemical studies are needed to clarify the molecular details of this interplay within the SOX4–USP10–HNRNPC–ubiquitin complex. Whether SOX4 similarly modulates the stability of other HNRNP family members or exerts analogous functions in other mesenchymal cell lineages warrants further investigation.

HNRNPC is a ubiquitously expressed member of the heterogeneous nuclear ribonucleoprotein family. RIP-seq revealed that transcripts recovered in HNRNPC immunoprecipitates are enriched in GO terms related to osteogenic differentiation and aortic development. We identified *WNT5A*, a prototypical member of the Wnt family that activates non-canonical Wnt signaling and has been implicated in osteochondrogenic differentiation and vascular calcification,^37^ as an HNRNPC-associated transcript whose stability depends on HNRNPC. Mechanistically, HNRNPC has been shown to stabilize target mRNAs by binding to their 3′-untranslated regions or coding sequences, thereby protecting them from exonucleolytic degradation.^31,32^ This binding can also be modulated by m⁶A modifications, which are recognized by HNRNPC and influence transcript stability.^36^ Based on these findings, we propose that HNRNPC similarly binds to and stabilizes *WNT5A* mRNA, thereby elevating WNT5A expression and driving VSMC osteogenic differentiation. Although the precise binding region on WNT5A mRNA remains to be mapped and a direct versus indirect mode of action is not resolved, our actinomycin D chase experiments establish that WNT5A mRNA stability is HNRNPC-dependent under both basal and pro-calcific conditions. Our data not only support the functional relevance of the SOX4–HNRNPC–WNT5A axis in governing VSMC fate, but also provide a mechanistic explanation for the previously reported inverse correlation between SOX4 and WNT5A expression.^51^ To directly validate the therapeutic potential of this pathway, we employed BOX5, a specific WNT5A antagonist. BOX5 treatment not only reversed the exacerbated calcification phenotype resulting from SOX4 deficiency but also potently suppressed osteogenic differentiation and calcification in wild-type VSMCs and mice under pro-calcific conditions. Therefore, these findings define the SOX4–HNRNPC–WNT5A axis as a regulatory mechanism in VSMC osteogenic differentiation and identify WNT5A, rather than SOX4, as the druggable node within it.

Our study has several limitations that merit discussion. First, the *Myh11*-CreERT2 mouse line used for VSMC-specific *Sox4* ablation is exclusively applicable to male mice; consequently, the potential influence of sex on the experimental results could not be assessed. Second, while we demonstrate that SOX4 regulates USP10-HNRNPC interaction, the precise ubiquitination sites and binding domain involved remain to be determined. Elucidating these molecular details will provide deeper mechanistic insight into SOX4-mediated regulation of HNRNPC stability. Third, our *in vivo* findings rely solely on the VitD3-induced vascular calcification model. Although this model is widely used and recapitulates key features of medial calcification, it does not fully capture the complexity of vascular calcification that occurs in the settings of chronic kidney disease, atherosclerosis, or diabetes. Future studies employing additional models will be essential to corroborate and extend our findings. Fourth, the relationship between HNRNPC and WNT5A mRNA is established here at the level of association and transcript stability; the binding site on WNT5A mRNA and the direct or indirect nature of this regulation remain to be determined.

## Conclusions

This study reveals that SOX4 restrains VSMC osteogenic differentiation by promoting HNRNPC-dependent destabilization of WNT5A mRNA. Targeting WNT5A with BOX5 attenuates VSMC osteogenic differentiation and vascular calcification in the vitamin D3–induced mouse model. However, WNT5A antagonism should be evaluated in disease-relevant models of calcific vasculopathy before translational conclusions are drawn.

## Acknowledgments

We thank Dr. She-long Zhang (Equipment and Technology Service Platform, College of Life Sciences, Zhejiang University) for their excellent technical support with microscopy.

## Author Contributions

C.Q., L.Y., and J.C. designed the project; J.C., H.C., X.J., Y.L., Y.Z., and K.M. conducted the experiments and data analysis; J.C. and C.Q. drafted the manuscript; H.L., X.Z, L.Y., and C.Q. revised the manuscript. All authors agree to be accountable for all aspects of work ensuring integrity and accuracy.

## References

1. Durham AL, Speer MY, Scatena M, Giachelli CM, Shanahan CM. Role of smooth muscle cells in vascular calcification: implications in atherosclerosis and arterial stiffness. Cardiovasc Res. 2018;114:590–600.

2. Lan Z, Liang Q, Li L, Liu F, Chen A, Ye Y, Feng L, Zhang Z, Zhang X, Ou JS. TRIM16 Mediates K63-Linked Ubiquitination of DAB2 to Facilitate Vascular Calcification. Circ Res. 2025;137:551–568.

3. Li W, Feng W, Su X, Luo D, Li Z, Zhou Y, Zhu Y, Zhang M, Chen J, Liu B, Huang H. SIRT6 protects vascular smooth muscle cells from osteogenic transdifferentiation via Runx2 in chronic kidney disease. J Clin Invest. 2022;132. doi:10.1172/JCI150051.

4. Leopold JA. Vascular calcification: Mechanisms of vascular smooth muscle cell calcification. Trends Cardiovasc Med. 2015;25:267–274.

5. Guo W, Guo W, Chen B, Lin Z, Wen Z, Ye J, Feng W, Feng X, Yan J, Yang P. NEXN protects against vascular calcification by promoting SERCA2 SUMOylation and stabilization. Nat Commun. 2025;16:8074.

6. Jin J, He Y, Guo J, Pan Q, Wei X, Xu C, Qi Z, Li Q, Ma S, Lin J. BACH1 controls hepatic insulin signaling and glucose homeostasis in mice. Nat Commun. 2023:8428.

7. Zhang X, Wang P, Liu LH, Zhang QT, Lv YQ, Yang HR, Hou YM, Li N, Cheng J, Wang ZY. Interleukin enhancer binding factor 3 exacerbates cardiac inflammation and injury following myocardial infarction by inhibiting Lys48-linked ubiquitination on HNRNPA2B1 in macrophages. Cell Mol Immunol. 2026:654–672.

8. Zhu L, Choudhary K, Gonzalez-Teran B, Ang YS, Thomas R, Stone NR, Liu L, Zhou P, Zhu C, Ruan H. Transcription Factor GATA4 Regulates Cell Type-Specific Splicing Through Direct Interaction With RNA in Human Induced Pluripotent Stem Cell-Derived Cardiac Progenitors. Circulation. 2022;146:770–787.

9. Tiwari N, Tiwari VK, Waldmeier L, Balwierz PJ, Arnold P, Pachkov M, Meyer-Schaller N, Schubeler D, van Nimwegen E, Christofori G. Sox4 is a master regulator of epithelial-mesenchymal transition by controlling Ezh2 expression and epigenetic reprogramming. Cancer Cell. 2013;23:768–783.

10. Cheng CK, Lin X, Pu Y, Tse JKY, Wang Y, Zhang CL, Cao X, Lau CW, Huang J, He L. SOX4 is a novel phenotypic regulator of endothelial cells in atherosclerosis revealed by single-cell analysis. J Adv Res. 2023;43:187–203.

11. Feng Y, Zhu N, Bedi K, Li J, Perera C, Green M, Assarzadegan N, Zhai Y, Liu Q, Baladandayuthapani V. SOX9 suppresses colon cancer via inhibiting epithelial-mesenchymal transition and SOX2 induction. J Clin Invest. 2025;135. doi:10.1172/JCI184115.

12. Fuglerud BM, Drissler S, Lotto J, Stephan TL, Thakur A, Cullum R, Hoodless PA. SOX9 reprograms endothelial cells by altering the chromatin landscape. Nucleic Acids Res. 2022;50:8547–8565.

13. Zhao J, Patel J, Kaur S, Sim SL, Wong HY, Styke C, Hogan I, Kahler S, Hamilton H, Wadlow R. Sox9 and Rbpj differentially regulate endothelial to mesenchymal transition and wound scarring in murine endovascular progenitors. Nat Commun. 2021;12:2564.

14. Huk DJ, Austin BF, Horne TE, Hinton RB, Ray WC, Heistad DD, Lincoln J. Valve Endothelial Cell-Derived Tgfbeta1 Signaling Promotes Nuclear Localization of Sox9 in Interstitial Cells Associated With Attenuated Calcification. Arterioscler Thromb Vasc Biol. 2016;36:328–338.

15. Faleeva M, Ahmad S, Theofilatos K, Lynham S, Watson G, Whitehead M, Marhuenda E, Iskratsch T, Cox S, Shanahan CM. Sox9 Accelerates Vascular Aging by Regulating Extracellular Matrix Composition and Stiffness. Circ Res. 2024;134:307–324.

16. Marom R, Song IW, Busse EC, Washington ME, Berrier AS, Rossi VC, Ortinau L, Jeong Y, Jiang MM, Dawson BC. The IFITM5 mutation in osteogenesis imperfecta type V is associated with an ERK/SOX9-dependent osteoprogenitor differentiation defect. J Clin Invest. 2024;134. doi:10.1172/JCI170369.

17. Angelozzi M, Karvande A, Lefebvre V. SOXC are critical regulators of adult bone mass. Nat Commun. 2024;15:2956.

18. Li Y, Ma C, Sheng Y, Huang S, Sun H, Ti Y, Wang Z, Wang F, Chen F, Li C. TRIB3 mediates vascular calcification by facilitating self-ubiquitination and dissociation of Smurf1 in chronic kidney disease. J Clin Invest. 2025;135. doi:10.1172/JCI175972.

19. Chiang HY, Chu PH, Chen SC, Lee TH. MFG-E8 promotes osteogenic transdifferentiation of smooth muscle cells and vascular calcification by regulating TGF-beta1 signaling. Commun Biol. 2022;5:364.

20. Xu L, Liu B, Ma H, Qi E, Ma J, Chang T, Zhang J, Zhang W, Chen W, Cao X, Xiong X. O-GlcNAc transferase promotes vascular smooth muscle calcification through modulating Wnt/beta-catenin signaling. FASEB J. 2024;38:e70271.

21. David CJ, Huang YH, Chen M, Su J, Zou Y, Bardeesy N, Iacobuzio-Donahue CA, Massague J. TGF-beta Tumor Suppression through a Lethal EMT. Cell. 2016;164:1015–1030.

22. Sun J, Xiong Y, Jiang K, Xin B, Jiang T, Wei R, Zou Y, Tan H, Jiang T, Yang A. Hypoxia-sensitive long noncoding RNA CASC15 promotes lung tumorigenesis by regulating the SOX4/beta-catenin axis. J Exp Clin Cancer Res. 2021;40:12.

23. Yang L, Dai R, Wu H, Cai Z, Xie N, Zhang X, Shen Y, Gong Z, Jia Y, Yu F. Unspliced XBP1 Counteracts beta-Catenin to Inhibit Vascular Calcification. Circ Res. 2022;130:213–229.

24. Xie Y, Lin T, Jin Y, Berezowitz AG, Wang XL, Lu J, Cai Y, Guzman RJ. Smooth muscle cell-specific matrix metalloproteinase 3 deletion reduces osteogenic transformation and medial artery calcification. Cardiovasc Res. 2024;120:658–670.

25. Harley VR, Clarkson MJ, Argentaro A. The molecular action and regulation of the testis-determining factors, SRY (sex-determining region on the Y chromosome) and SOX9 [SRY-related high-mobility group (HMG) box 9]. Endocr Rev. 2003;24:466–487.

26. Wang G, Xiao Q, Luo Z, Ye S, Xu Q. Functional impact of heterogeneous nuclear ribonucleoprotein A2/B1 in smooth muscle differentiation from stem cells and embryonic arteriogenesis. J Biol Chem. 2012;287:2896–2906.

27. Chen Q, Jin M, Zhu J, Xiao Q, Zhang L. Functions of heterogeneous nuclear ribonucleoproteins in stem cell potency and differentiation. Biomed Res Int. 2013;2013:623978.

28. Wang Y, Yang T, Li Q, Zheng Z, Liao L, Cen J, Chen W, Luo J, Xu Y, Zhou M, Zhang J. circASAP1 induces renal clear cell carcinoma ferroptosis by binding to HNRNPC and thereby regulating GPX4. Mol Cancer. 2025;24:1.

29. Xia X, Liu X, Chai R, Xu Q, Luo Z, Gu J, Jin Y, Hu T, Yu C, Du B. USP10 exacerbates neointima formation by stabilizing Skp2 protein in vascular smooth muscle cells. J Biol Chem. 2021;297:101258.

30. Zhang W, Zhao J, Deng L, Ishimwe N, Pauli J, Wu W, Shan S, Kempf W, Ballantyne MD, Kim D. INKILN is a Novel Long Noncoding RNA Promoting Vascular Smooth Muscle Inflammation via Scaffolding MKL1 and USP10. Circulation. 2023;148:47–67.

31. Huang L, Yang G, Shao Y, Sun J, Yang X, Hong H, Aikemu B, Yesseyeva G, Li S, Ding C. Cancer-derived exosomal lncRNA SNHG3 promotes the metastasis of colorectal cancer through hnRNPC-mediating RNA stability of beta-catenin. Int J Biol Sci. 2024;20:2388–2402.

32. Chen JJ, Lu TZ, Wang T, Yan WH, Zhong FY, Qu XH, Gong XC, Li JG, Tou FF, Jiang LP, Han XJ. The m6A reader HNRNPC promotes glioma progression by enhancing the stability of IRAK1 mRNA through the MAPK pathway. Cell Death Dis. 2024;15:390.

33. Xiong X, Feng S, Ma X, Liu K, Gui Y, Chen B, Fan X, Wang F, Wang X, Yuan S. hnRNPC Functions with HuR to Regulate Alternative Splicing in an m6A-Dependent Manner and is Essential for Meiosis. Adv Sci (Weinh). 2025;12:e2412196.

34. Zheng Z, Zeng X, Zhu Y, Leng M, Zhang Z, Wang Q, Liu X, Zeng S, Xiao Y, Hu C. CircPPAP2B controls metastasis of clear cell renal cell carcinoma via HNRNPC-dependent alternative splicing and targeting the miR-182-5p/CYP1B1 axis. Mol Cancer. 2024;23:4.

35. Liu YY, Xia M, Chen ZB, Liao YD, Zhang CY, Yuan L, Pan YW, Huang H, Lu HW, Yao SZ. HNRNPC mediates lymphatic metastasis of cervical cancer through m6A-dependent alternative splicing of FOXM1. Cell Death Dis. 2024;15:732.

36. De Kesel J, Fijalkowski I, Pieters T, Borin C, Christensen KT, Wittouck M, Van Laere J, Guerrero L, Reunes L, Caruso M. HNRNPC and m6A RNA methylation control oncogenic transcription and metabolism in T-cell leukemia. Blood. 2025;146:275–290.

37. Woldt E, Terrand J, Mlih M, Matz RL, Bruban V, Coudane F, Foppolo S, El Asmar Z, Chollet ME, Ninio E. The nuclear hormone receptor PPARgamma counteracts vascular calcification by inhibiting Wnt5a signalling in vascular smooth muscle cells. Nat Commun. 2012;3:1077.

38. Kawarazaki W, Mizuno R, Nishimoto M, Ayuzawa N, Hirohama D, Ueda K, Kawakami-Mori F, Oba S, Marumo T, Fujita T. Salt causes aging-associated hypertension via vascular Wnt5a under Klotho deficiency. J Clin Invest. 2020;130:4152–4166.

39. Gu S, Feng H, Li X, Dong Y, Peng Y, Liu Q, Zhang X, You J, Zhu J, Zhao X. Targeting WNT5A noncanonical signaling attenuates renal fibrosis progression in acute kidney injury. Mol Ther. 2025;33:5248–5262.

40. Liu D, Du J, Xie H, Tian H, Lu L, Zhang C, Xu GT, Zhang J. Wnt5a/beta-catenin-mediated epithelial-mesenchymal transition: a key driver of subretinal fibrosis in neovascular age-related macular degeneration. J Neuroinflammation. 2024;21:75.

41. Wu J, Xu H, Wang S, Weng H, Luo Z, Ou G, Chen Y, Xu L, So KF, Deng L. Regular exercise ameliorates high-fat diet-induced depressive-like behaviors by activating hippocampal neuronal autophagy and enhancing synaptic plasticity. Cell Death Dis. 2024;15:737.

42. Lanzer P, Hannan FM, Lanzer JD, Janzen J, Raggi P, Furniss D, Schuchardt M, Thakker R, Fok PW, Saez-Rodriguez J. Medial Arterial Calcification: JACC State-of-the-Art Review. J Am Coll Cardiol. 2021;78:1145–1165.

43. St Hilaire C. Medial Arterial Calcification: A Significant and Independent Contributor of Peripheral Artery Disease. Arterioscler Thromb Vasc Biol. 2022;42:253–260.

44. Zhang H, Li G, Yu X, Yang J, Jiang A, Cheng H, Fu J, Liang X, Liu J, Lou J. Progression of Vascular Calcification and Clinical Outcomes in Patients Receiving Maintenance Dialysis. JAMA Netw Open. 2023;6:e2310909.

45. Bin S, Xinyi F, Huan P, Xiaoqin Z, Jiming W, Yi H, Ziyue L, Xiaochun Z, Zhouqi L, Bangwei Z. SOX4 as a potential therapeutic target for pathological cardiac hypertrophy. Eur J Pharmacol. 2023;958:176071.

46. Zhang L, Lv L, Zheng N, Li R, Yang R, Li T, Li Y, Liu Y, Luo H, Li X. Suppression of Sox4 protects against myocardial ischemic injury by reduction of cardiac apoptosis in mice. J Cell Physiol. 2021;236:1094–1104.

47. Dong Q, Lu L, Zhang X, Feng L, Li L, Liu H, Chen A, Lan Z, Ye Y, Zhu J. NAMPT Is A Novel Inhibitor of Vascular Calcification in Chronic Kidney Disease. Arterioscler Thromb Vasc Biol. 2025;45:1872–1892.

48. Willems BA, Furmanik M, Caron MMJ, Chatrou MLL, Kusters DHM, Welting TJM, Stock M, Rafael MS, Viegas CSB, Simes DC. Ucma/GRP inhibits phosphate-induced vascular smooth muscle cell calcification via SMAD-dependent BMP signalling. Sci Rep. 2018;8:4961.

49. Zhang M, Wu H, Lin X, Zhao F, Deng Q, Lu L, Li X, Chen X, Liu H, Love PE. MYC-induced USP10 stabilizes SOX4 to promote thymocyte proliferation and leukemia onset in mice. Nat Commun. 2026. doi:10.1038/s41467-026-71084-w.

50. Pan X, Zhao J, Zhang WN, Li HY, Mu R, Zhou T, Zhang HY, Gong WL, Yu M, Man JH. Induction of SOX4 by DNA damage is critical for p53 stabilization and function. Proc Natl Acad Sci U S A. 2009;106:3788–3793.

51. Moran JD, Kim HH, Li Z, Moreno CS. SOX4 regulates invasion of bladder cancer cells via repression of WNT5a. Int J Oncol. 2019;55:359–370.

